# Brain-Environment Alignment during Movie Watching Predicts Cognitive-Affective Function in Adulthood

**DOI:** 10.1101/2020.09.15.298125

**Authors:** Raluca Petrican, Kim S. Graham, Andrew D. Lawrence

## Abstract

BOLD fMRI studies have provided compelling evidence that the human brain demonstrates substantial moment-to-moment fluctuations in both activity and functional connectivity patterns. While the role of brain signal variability in fostering cognitive adaptation to ongoing environmental demands is well-documented, the relevance of moment-to-moment changes in functional brain architecture is still debated. To probe the role of architectural variability in naturalistic information processing, we used neuroimaging and behavioural data collected during movie watching by the Cambridge Centre for Ageing and Neuroscience (N = 642, 326 women) and the Human Connectome Project (N = 176, 106 women). Both moment-to-moment and contextual change-evoked architectural variability increased from young to older adulthood. However, coupling between moment-to-moment changes in functional brain architecture and concrete environmental features was stronger at younger ages. Architectural variability (both momentary and context-evoked) was associated with age-distinct profiles of network communication, specifically, greater functional integration of the default mode network in older adulthood, but greater informational flow across neural networks implicated in environmentally driven attention and control (cingulo-opercular, salience, ventral attention) in younger adulthood. Whole-brain communication pathways anchored in default mode regions relevant to episodic and semantic context creation (i.e., angular and middle temporal gyri) contributed to greater brain reconfiguration in response to narrative context changes, as well as stronger coupling between moment-to-moment changes in functional brain architecture and changes in concrete environmental features. Cognitive adaptation was directly linked to levels of brain-environment alignment, but only indirectly associated with levels of architectural variability. Specifically, stronger coupling between moment-to-moment variability in brain architecture and concrete environmental features predicted poorer cognitive adaptation (i.e., fluid IQ) and greater affectively driven environmental vigilance. Complementarily, across the adult lifespan, higher fluid (but not crystallized) IQ was related to stronger expression of the network communication profile underlying momentary and context-based architectural variability during youth. Our results indicate that the adaptiveness of dynamic brain reconfiguration during naturalistic information processing changes across the lifespan due to the associated network communication profiles. Moreover, our findings on brain-environment alignment complement the existing literature on the beneficial consequences of modulating brain signal variability in response to environmental complexity. Specifically, they imply that coupling between moment-to-moment variability in functional brain architecture and concrete environmental features may index a bias towards perceptually-bound, rather than conceptual processing, which hinders affective functioning and strategic engagement with the external environment.

---

BOLD fMRI studies testify to the importance of acknowledging the highly dynamic nature of the human brain in order to better understand its lifespan developmental trajectory (Garrett et al., 2015, 2017, 2020; Grady & Garrett, 2014; Poldrack & Shine, 2018). A substantial portion of this research has focused on moment-to-moment brain signal variability and provided compelling evidence that its flexible modulation based on current task requirements (e.g., external vs. internal attentional focus, difficulty level) is key to optimal functioning (Garrett et al., 2020; Grady & Garrett, 2018; Roberts, Grady, & Addis, 2020). Conversely, age-related cognitive deficits are thought to stem from age-related decay in the ability to adjust brain signal variability in response to ongoing demands (Garrett, Kovacevic, McIntosh, & Grady, 2013a). A consensus emerging from this literature is that brain signal variability is likely to provide the basis for neural flexibility, underpinning the capacity of a brain to tune into the dynamics of the external world and respond in a differentiated manner to a wide range of environmental stimuli (Garrett et al., 2020; Grady & Garrett, 2018; Padmanabhan & Urban, 2010).

Importantly, the human brain demonstrates substantial moment-to-moment fluctuations not only in activity, but also in functional connectivity patterns (Poldrack & Shine, 2018). However, the relevance of architectural variability is still debated, with some studies underscoring its role in maturation and learning, whereas others point to its association with accelerated cognitive ageing in later life (Bassett et al., 2011, 2013; Hutchison & Morton, 2015; Mujica-Parodi et al., 2020).

Here, we seek to probe this issue further by testing whether, similar to signal variability, fluctuations in functional neural architecture can facilitate alignment of brain-environment dynamics and, thus, contribute to flexible responding to the external world. To this end, we estimate patterns of whole-brain functional reconfiguration during movie watching, a dynamic cognition paradigm well-suited for characterising individual and ageing-related differences in real-life information processing (Bottenhorn et al., 2019; Demirtas et al., 2019; Geerligs, Cam-Can, & Campbell, 2018; Simony et al., 2016; Sonkusare, Breakspear, & Guo, 2019). We thus capitalise on evidence that the human mind readily breaks down the continuous influx of environmental information into meaningful units, which it uses to predict and encode ongoing experiences (Baldwin & Kosie, 2020; Kurby & Zacks, 2008). Such segmentation processes unfold at multiple timescales and give rise to a nested event hierarchy, which spans relatively frequent and readily processed low-level featural fluctuations (e.g., object presence/absence) to more sporadic and abstract, slower processed, high-level changes to the “working event model” underlying an ongoing experience (Baldassano et al., 2018; Zacks, 2020).

Applying insights from the literature on brain signal variability, here, we investigate whether neural architectural variability associated with (1) moment-to-moment fluctuations in concrete environmental features and (2) changes to the more abstract “working event model” could also be a reflection of a brain’s dynamic range, specifically, of its capacity to respond in a more differentiated manner and, thus, better adapt to the external world (McDonnell & Ward, 2011). To this end, we tested whether either type of architectural variability predicts fluid intelligence, a mental capacity reliant on well-differentiated perceptual representations of the external world, which supports flexible updating of event representations and successful adaptation to novel environments (Cattell, 1971; Colzato, van Wouwe, Lavender, & Hommel, 2006; Duncan, Assem, & Shashidhara, 2020; Duncan, Chylinskia, Mitchell, & Bhandaric, 2017). To probe the specificity of any observed effects, we included a crystallized intelligence measure, which, as an index of accumulated knowledge, was not expected to be linked to our presumed indices of neural flexibility (i.e., the two types of architectural variability).

Brain signal variability has been shown to decline with ageing and, thus, contribute to some of the cognitive deficits observed in older adulthood (Garrett, Kovacevic, McIntosh, & Grady, 2010; Garrett et al., 2020). Consequently, our second objective was to establish whether levels of moment-to-moment and context-based architectural variability also fluctuate across the adult lifespan and whether any such fluctuations are linked to age-related differences in fluid intelligence. If either type of architectural variability reflects neural flexibility, then we would expect it to decline in older adulthood, just like brain signal variability does (Garrett et al., 2010, 2013a), with its decline predicting fluid intelligence decrements. If, on the other hand, greater functional brain reconfiguration during movie watching is an index of architectural instability and, thus, deficient neural functioning, as it has been reported for resting state and experimental task paradigms (cf. Mujica-Parodi et al., 2020), then we would expect it to increase with age, with increments in variability predicting decrements in fluid intelligence.

Beyond mean variability levels, environmentally-driven modulation of brain signal variability, presumably a more sensitive index of context-based neural differentiation, has been uniquely linked to superior cognitive functioning (Garrett et al., 2020). Consequently, our third objective was to probe whether strength of coupling between neural architectural changes and changes in concrete environmental features would predict fluid intelligence beyond mean levels of moment-to-moment architectural variability. To gauge the affective relevance of alignment in brain organisation-concrete environment dynamics, we included depression and anxiety scales because the two capture complementary affectively-driven biases in attention (i.e., towards abstract, internally generated versus concrete, external information), which shape naturalistic information processing (Belzung, Wilner, & Philippot, 2015; Brewin, Gregory, Lipton, & Burgess, 2010; Hermans, Henckens, Joels, & Fernandez, 2014; Sherrill, Kurby, Lilly, & Magliano, 2019; Sylverster et al., 2012). If greater coupling between changes in brain architecture and concrete environmental features reflects stronger neural tuning to the dynamics of the external world, then we would expect it to be positively linked to anxiety, but negatively linked to depression (Petrican, Saverino, Rosenbaum, & Grady, 2015).

Studies on brain signal variability underscore the adaptiveness of activity fluctuations in functionally specific regions and networks, most widely, those involved in attending to the external world (visual, salience [SAL], dorsal attention [DAN], Garrett et al., 2020; Grady & Garrett, 2018). Similarly, literature on adaptive architectural variability supporting learning highlights its association with greater functional integration of systems involved in external attention and environmentally driven control (SAL, DAN, cingulo-opercular [CON]), but weaker integration of these systems with the network generally implicated in self-generated cognition (default mode network, [DMN]) (Finc et al., 2020). These findings thus raise the possibility that it may not be levels of architectural variability per se, but rather their associated network communication profiles which determine the adaptiveness of functional brain reconfiguration during real-life information processing. Hence, our fourth objective was to test whether moment-to-moment and context-based architectural variability are associated with distinct patterns of network integration across the adult lifespan and whether it is these age-distinct network profiles, rather than variability levels, that predict fluid intelligence. With age, modulation of brain signal variability in response to external environmental stimuli declines, while the functional dominance of systems implicated in internal cognition increases (Garrett et al., 2013a; Grady & Garrett, 2018; Spreng & Turner, 2019). Consequently, we anticipated that greater architectural variability would be associated with greater informational flow (i.e., integration) over systems involved in external processing (CON, SAL, DAN, ventral attention [VAN], perceptual systems) at younger ages, but greater functional integration of the DMN in older adulthood. As an exploratory analysis, we further sought to link event perception processes across multiple timescales by characterising brain region-specific patterns of informational flow which support brain-environment alignment with respect to both contextual and concrete featural changes.

To gauge variability in functional brain architecture, we adopted a graph theoretical approach (see Method for details), similar to the one previously used to probe the adaptiveness of dynamic neural reconfiguration during learning (Bassett et al., 2011, 2013). An advantage of this approach is that architectural variability is defined at the level of the whole-brain for each individual separately. This means that changes in specific neural connections are “contextualized” with respect to how the remaining connections change, an aspect which we deemed important for enhancing the sensitivity of our analyses given prior evidence of idiosyncrasies in brain organisation (cf. Gordon et al., 2017).

The present report is structured as follows. Part 1 focuses on a healthy adult lifespan sample from the Cambridge Centre for Ageing and Neuroscience. Its purpose is to investigate whether (1) levels of moment-to-moment versus context-based architectural variability fluctuate across the adult lifespan, whether (2) they are associated with age-distinct profiles of network integration (i.e., informational flow) and whether (3) any such age differences in levels of architectural variability or in their associated network profiles predict differences in fluid intelligence. Part 2 is based on data from a separate sample of healthy young adult participants in the Human Connectome Project. Its aim is two-fold. The first is to probe the cognitive (i.e., fluid intelligence) and affective (i.e., depression/anxiety) relevance of alignment in neural architectural fluctuations and changes in concrete environmental features. The second is to link event segmentation processes across multiple time scales by characterising patterns of brain region-specific informational flow underlying dynamic brain reconfiguration in response to both abstract contextual changes and concrete featural fluctuations.

## Part 1: Cam-Can Sample

### Methods

#### Participants

We included the largest number of participants from Stage II of the Cambridge Ageing and Neuroscience (Cam-Can) study with available fMRI data in the movie watching condition (N = 642, age range: 18-88 yrs [M = 54 yrs, SD = 19 yrs]).

The majority of participants (N = 589) were predominantly right-handed (handedness measure > 0). The sample included 316 men (32 between 18 and 29, 49 between 30 and 39, 48 between 40 and 49, 48 between 50 and 59, 57 between 60 and 69, 50 between 70 and 79, 32 between 80 and 88 years of age) and 326 women (42 between 18 and 29, 46 between 30 and 39, 57 between 40 and 49, 45 between 50 and 59, 48 between 60 and 69, 58 between 70 and 79, 30 between 80 and 88 years of age).

All participants were cognitively healthy (MMSE > 24) and met the hearing, vision, and English language ability criteria necessary for the completing experimental tasks (Taylor et al., 2017). They were also screened for any neurological and serious psychiatric conditions, as well as for physical conditions or bodily implants that may render their participation unsafe (Taylor et al., 2017). Participants provided informed consent in accordance with the Cambridgeshire research ethics board (Shafto et al., 2014).

#### Out-of-scanner measures

##### Fluid intelligence

Participants completed the pen-and-paper version of the Cattell Culture Fair Test, Scale 2 Form A (Cattell, 1971; Cattell & Cattell, 1973) in which they had to select the correct answer from multiple alternatives and record it on an answer sheet. The test contains four subtests with distinct non-verbal “puzzles”: series completion, classification, matrices and conditions. Unbeknownst to the participants, each subtest is timed, such that 3, 4, 3 and 2.5 minutes are allocated for subtest 1, 2, 3 and 4, respectively. Correct responses receive a score of 1 for a maximum score of 46.

##### Crystallized intelligence

The proverb comprehension test was used as an index of crystalized intelligence (Crawford & Stankov, 1996). In this task, participants provide the meaning of three common proverbs in English, which are presented on a computer screen (Shafto et al., 2014). Their responses are recorded digitally and scored by experimenters as incorrect or a “don’t know” response (0), partly correct but literal rather than abstract (1), or fully correct and abstract (2). The highest possible score is 6.

#### In-scanner task

Participants watched a shortened version of Alfred Hitchcock’s black-and-white television drama “Bang! You’re Dead” (Hitchcock, 1961; Hasson et al., 2008, 2010) edited from 30 min to 8 min while maintaining the plot (Shafto et al., 2014). The movie was selected to be engaging and unfamiliar to the participants.

#### fMRI data acquisition

Images were acquired with a 3T Siemens TIM Trio System (32-channel coil). T1-weighted anatomical scans were acquired with a MPRAGE sequence (TR = 2250 ms, TE = 2.99 ms, flip angle=9°, FOV = 256 mm x 240 mm x 192 mm, 1 mm isotropic voxels, GRAPPA acceleration factor = 2). Functional images were acquired with a multi-echo EPI sequence (TR=2470 ms, [TE=9.4 ms, 21.2 ms, 33 ms, 45 ms, 57 ms], flip angle=78°, FOV = 192 x 192 mm, 32 axial slices of 3.7 mm thickness, acquired in descending order with 20% gap, voxel size of 3 mm × 3 mm x 4.4 mm).

#### fMRI data preprocessing

The main preprocessing and analysis steps for both samples are presented in Figure 1. For the Cam-Can data, preprocessing began with the averaging of the corresponding images from the multiple echos. We opted for a simple average, rather than a weighted sum, in order to maximize comparability with the HCP data, which did not contain multiple echos. We reasoned that our strategy was defensible in light of evidence that simple averaging yields similar improvements in 3T image quality as weighted summing (Kettinger et al., 2016; Poser, Versluis, Hoogduin, & Norris, 2006). Subsequently, we performed image processing in SPM12 (Wellcome Department of Imaging Neuroscience, London, UK). Specifically, we corrected for slice timing differences and rigid body motion (which included unwarping), spatially normalized the images to the standard Montreal Neurological Institute (MNI)-152 template, and smoothed them (full-width half-maximum, 6 mm).

**Figure 1.**
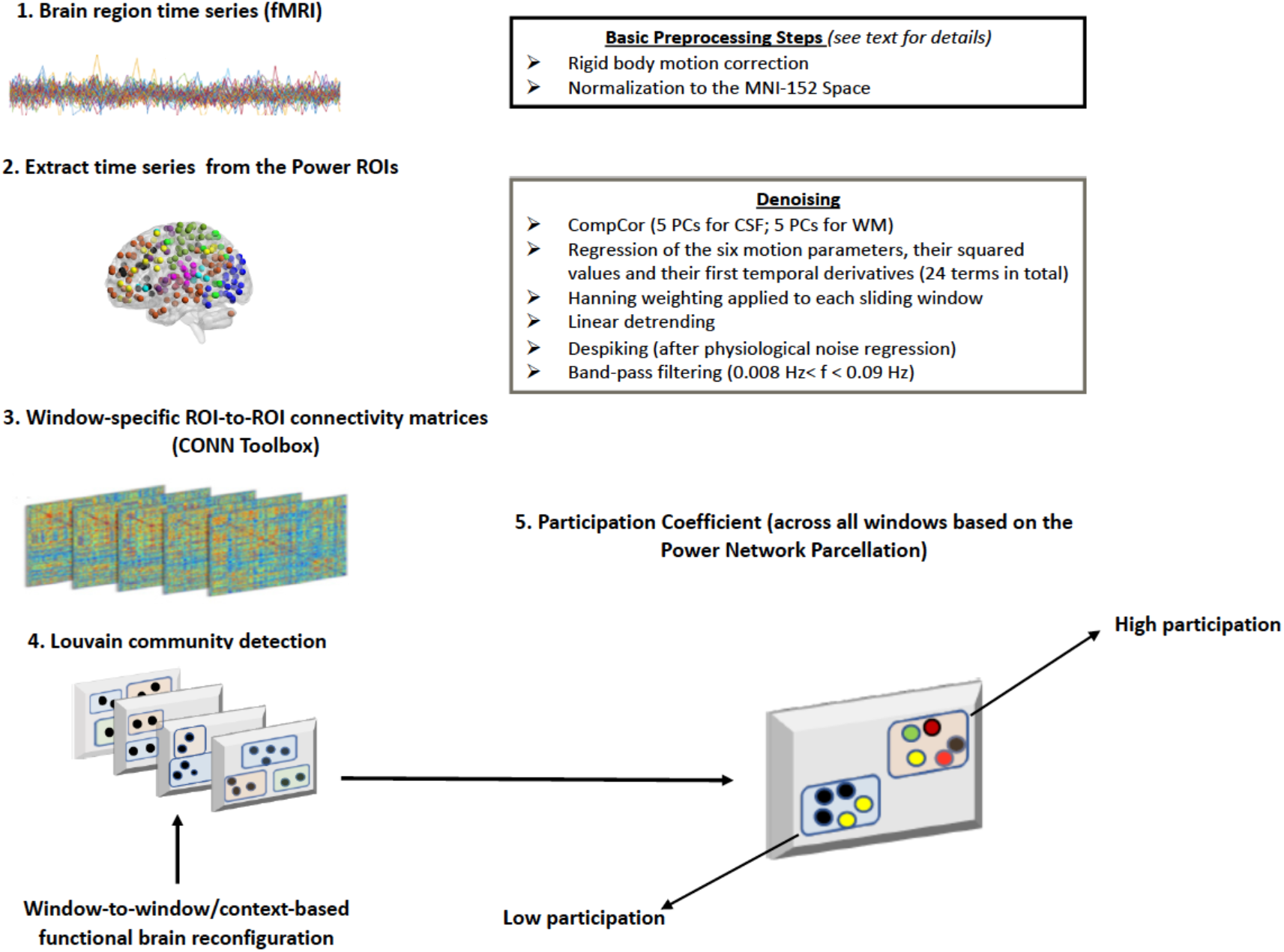
Schematic representation of the main preprocessing and analysis steps across both samples.

##### Additional denoising

Because motion can significantly impact functional connectivity measures (Power et al., 2012; Van Dijk et al., 2012), we implemented several additional preprocessing steps to address this potential confound. First, after extracting the BOLD time series from our regions-of-interest (ROIs, see below), but prior to computing the ROI-to-ROI correlations, we used the Denoising step in the CONN toolbox (version 17c; Whitfield-Gabrieli & Nieto-Castanon, 2012) to apply further physiological and rigid motion corrections. Specifically, linear regression was used to remove from the BOLD time series of each ROI the BOLD time series of the voxels within the MNI-152 white matter and CSF masks, respectively (i.e., the default CONN option of five CompCor-extracted principal components for each, Behzadi, Restom, Liau, & Liu, 2007), the 6 realignment parameters, their first-order temporal derivatives and their associated quadratic terms (24 regressors in total, cf. Bolt et al., 2017). The residual BOLD time series were bandpass filtered (0.008 Hz< f < 0.09 Hz), linearly detrended and despiked (all three are default CONN denoising steps). Following these corrections (which did not include global signal regression) (Murphy & Fox, 2017), an inspection of each subject’s histogram of voxel-to-voxel connectivity values revealed a normal distribution, approximately centered around zero, which would suggest reduced contamination from physiological and motion-related confounds (cf. Whietfield-Gabrieli & Castanon, 2012). Nonetheless, in all hypothesis testing analyses, we controlled for the average relative (i.e., volume-to-volume) displacement per participant, a widely used motion metric (Power et al., 2012, 2015; Satterthwaite et al., 2013).

#### fMRI data analysis

##### ROI time series

229 nodes for 10 core large-scale functional brain networks (i.e., default-mode [DMN], frontoparietal [FPC], cingulo-opercular [CON], salience [SAL], dorsal attention [DAN], ventral attention [VAN], somatomotor [SM], subcortical [SUB], auditory [AUD] and visual [VIS]) were defined for each participant as spherical ROIs (radius 5 mm) centered on the coordinates of the regions reported in Power et al. (2011) and assigned network labels corresponding to the graph analyses from this earlier article. We selected the Power et al. atlas because it was created by taking into account both the task-related activation (derived meta-analytically) and the resting state connectivity patterns of the component voxels for each ROI. Thus, this atlas provided an optimal parcellation scheme for charactering functional brain reorganization during a naturalistic cognition condition that, due to the lack of an explicit task, was likely to share significant similarities with a resting state condition (Vanderwal et al., 2017).

The ROIs were created in FSL (Smith et al., 2004), using its standard 2 mm isotropic space, with each ROI containing 81 voxels. These template space dimensions were selected because they yielded the most adequate spatial representation of the Power atlas. The 229 ROIs represent a subset of the 264 putative functional areas proposed by Power et al. (2011). The 229 ROIs were selected because, based on Power et al.’s analyses, they showed relatively unambiguous membership to one of the ten large-scale functional networks outlined above.

###### Fit of the Power atlas to the present dataset

To evaluate the homogeneity of the Power ROIs across the Cam-Can adult lifespan sample (cf. Iraji et al., 2020), we used an approach that is conceptually similar to those recently used in the literature (Gordon et al., 2016; Siegel et al., 2016). Specifically, we used the CONN toolbox to compute the radial similarity contrast (RSC) for each voxel in the Power atlas. As implemented in CONN, the RSC reflects the amount of similarity in whole brain connectivity patterns between a voxel and its neighbours in each of the three space directions (x,y,z), and is thus a 3 dimensional construct (see also Kim et al., 2010). If a node is functionally homogenous, then the RSC of its voxels should be relatively similar (i.e., across the entire ROI, there should be a consistent degree of similarity among the component voxels’ whole brain connectivity patterns). To test this hypothesis, for each participant, we conducted a principal components analysis of the RSC of the 81 voxels within each ROI (i.e., for each participant, one ROI constituted one case, whereas the RSC of one voxel within a given ROI constituted a variable). As a measure of similarity among the RSC of all voxels within a node across all nodes, we took the percent of variance explained by the first component extracted through the analysis just described (see also Gordon et al., 2016; Siegel et al., 2016). Because, as previously mentioned, RSC is a three-dimensional vector, this set of analyses yielded, for each participant, three indices of average similarity in global functional connectivity patterns among all voxels within a given ROI. The three indices were significantly positively correlated across participants (Spearman’s *rhos* from .17 to .24, all *ps* < .0001), which is why we averaged them to create a summary measure of ROI functional homogeneity. A correlational analysis, based on 100,000 permutation samples, revealed a modest, but significant negative relationship between age and the average similarity in global functional connectivity patterns among the component voxels of the Power ROIs (Spearman’s *rho* of −.21, *p* = 10^-5^). As expected, given that the Power ROIs were validated on a sample corresponding in age to the first decade of the Cam-Can sample, a second robust regression analysis revealed no evidence of significant quadratic age effects on the functional homogeneity of the Power ROIs (*p* > 0.56). Based on these results, the summary measure of ROI functional homogeneity was introduced as a covariate in all hypothesis testing analyses.

###### Fit of the Power network assignment to the present dataset

To estimate the fit of Power et al.’s (2011) network assignments to the present data, we used the same algorithm utilized by these authors for community detection, i.e., Infomap (Lancichinetti & Fortunato, 2009; Rosvall & Berstrom, 2008). We ran our analyses using Infomap Online (D. Edler & M. Rosvall) available at www.mapequation.org/infomap.

Following Power et al. (2011), we conducted all our community detection analyses on group-averaged ROI-to-ROI correlation matrices. To test whether Power et al.’s (2011) network assignments provide an equally adequate fit to the present connectivity data across the lifespan, we first averaged the individuals’ ROI-to-ROI correlation matrices, obtained from CONN, separately within each of the seven decades. Next, we used the Brain Connectivity Toolbox (BCT, Rubinov & Sporns, 2010) to threshold each of the seven matrices at the same tie densities used by Power et al. (2011) for the areal graph (i.e., 2% to 10% tie density in increments of 1%).

BCT was subsequently utilized to write the thresholded matrices into a Pajek *.net format and the resulting files were inputted into Infomap. Infomap was run with the option of extracting a one-level structure (i.e., no nested modules), which is the format of Power et al.’s (2011) atlas as is publicly available. Finally, we used again the BCT to compute the normalized mutual information index (NMI; range: 1 [perfect similarity] to 0 [no similarity at all]) as a measure of similarity between Power et al.’s network assignments and those outputted by Infomap for the connectivity data from each of the Cam-Can’s seven decades (Fornito, Zalesky & Bullmore, 2016). As can be seen in Figure 2, across all tie densities, the network structure from each of the seven age groups exhibited equivalent levels of similarity with that proposed by Power et al. (2011). These results thus suggest that Power et al.’s network structure provide an adequate fit to the present connectivity data across the lifespan (i.e., the NMI values were similar to those documented by Power et al. when comparing network structure across cohorts).

**Figure 2.**
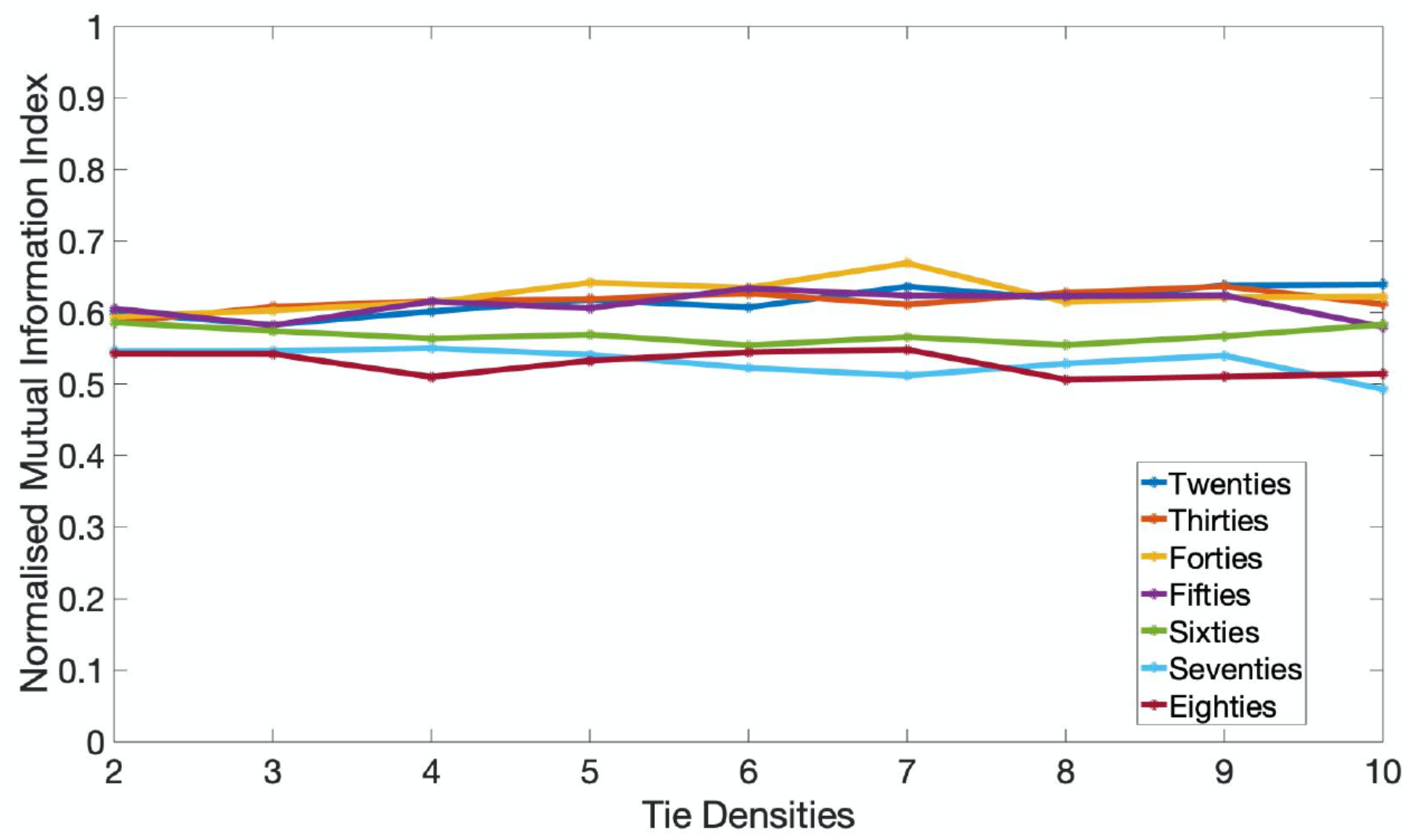
Network structure in each of the seven age groups of the Cam-Can sample during movie watching at 2-10% tie densities shows similar levels of similarity to that reported by Power et al.’ (2011) across various cohorts during rest. The colored lines show the normalized mutual information index (NMI), a highly used metric of similarity in community assignment, for each age group and tie density scrutinized.

##### Functional connectivity analyses

Pairwise coupling among the 229 ROIs was estimated in CONN. Pairwise correlations among all the ROIs were expressed as Fisher’s z-scores. Consistent with existing practices aimed at maximizing interpretability of results in network neuroscience studies of individual or group differences (e.g., sex or age, Betzel et al., 2014; Satterthwaite et al., 2015), we used both positive and negative z-scores to compute the indices of interest for all connectivity analyses. We reasoned that such an approach would be particularly well-justified in our present case since global signal regression, an artefact removal technique that generates negative correlations whose interpretation is still controversial, was not part of our preprocessing pipeline (for further discussion on the validity of the negative correlations obtained with the CONN toolbox, see Whitfield-Gabrieli & Nieto-Castanon, 2012).

To characterize individual differences in dynamic network structure, we used a combined window and clustering based approach (Iraji et al., 2020; Lurie et al., 2020). Thus, we broke down the movie into partially overlapping 40 s long windows for a total of 177 windows. This window length was selected in light of prior evidence that it both maximizes detection of individual differences in dynamic network reconfiguration and enables identification of a stable functional core (Leonardi & Van De Ville, 2015; Preti, Bolton, de Ville, 2017; Telesford et al., 2016; for similar window sizes in dynamic connectivity analyses of HCP data, see also Chen et al., 2016). Thus, pairwise coupling among the 229 ROIs was estimated in CONN using a sliding window of 40 s in length (~ 16 volumes), with one TR (2.47 s) gap and a “hanning weighting” (i.e., greater weight to the scans in the middle of the window relative to the ones at the periphery) applied to all the time points within a window. The use of a hanning weighting was intended to reduce the autocorrelation in the fMRI data series and, thus, maximize the opportunity to detect differences in functional brain organization between adjacent windows.

##### Network-level analyses

All the network-level metrics were computed using the Brain Connectivity Toolbox (BCT, Rubinov & Sporns, 2010) and the Network Community Toolbox (Bassett, D.S. [2017, November]. Network Community Toolbox. Retrieved from http://commdetect.weebly.com/), as described below.

##### Community detection

Rather than being computed directly, the degree to which a network can be fragmented into well-delineated and non-overlapping communities or modules is estimated using optimization algorithms, which sacrifice some degree of accuracy for processing speed (Fornito et al., 2016; Rubinov & Sporns, 2010). Here, the optimal whole-brain division into constituent communities was estimated using a Louvain community detection algorithm implemented in the BCT. This algorithm partitions a network into non-overlapping groups of nodes with the goal of maximizing an objective modularity quality function, Q (Betzel & Bassett, 2017; Rubinov & Sporns, 2011; Sporns & Betzel, 2016). There are multiple strategies for estimating community structure based on sliding window data, as was the case of our movie viewing data. Specifically, multilayer modularity algorithms (Bassett et al., 2011; Braun et al., 2015; Mucha et al., 2010) can provide important insights into community dynamics at multiple time scales. Nonetheless, such algorithms require estimation of additional free parameters (e.g., the temporal coupling parameter between two adjacent temporal windows). Since we feared that estimation of the temporal coupling parameter could act as a potential confound when comparing connectivity results across the multiple samples included in the analysis, we used the same procedure to estimate community structure independently in each sliding window (see also Chen et al., 2016), as described below.

For signed networks, such as the ones investigated in our study, optimization of the Q function can be achieved by either placing equal weight on maximizing positive within-module connections and minimizing negative within-module connections or by putting a premium on maximizing positive connections, which have been argued to be of greater biological significance (Fornito et al., 2016; Rubinov & Sporns, 2011). Although we verified that all the reported results emerge with either formula, for the sake of simplicity and because we agree with their argument regarding the greater importance of positive weights in determining node grouping into communities, we report here the results based on Rubinov and Sporns’s modularity formula (cf. Chen et al., 2016; Rubinov & Sporns, 2011). In this formulation, the contribution of positive weights to Q is not affected by the presence of negative weights in the network, whereas the contribution of negative weights to Q decreases with an increase in positive weights.

The adapted modularity function Q*, proposed by Rubinov and Sporns (2011) is written
as

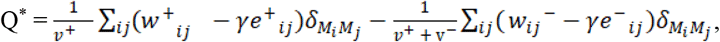

where δ_MiMj_ = 1 if nodes i and j are in the same module and δ_MiMj_ = 0 otherwise; v^+^ and v^-^ constitute the sum of all positive (w^+^) and all negative (w^-^) weights in the network, respectively; w^±^_ij_ represent the actual within-module positive or negative connection weights with w^±^ ∈ (0,1]; γ is a resolution parameter determining the size of the identified modules; e^±^_ij_ is the within-module connection strength expected by chance and defined, for each node-to-node (i,j) connection as 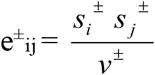, with s^±^_i_ and s^±^_j_ being the sum of all positive or all negative connection weights of node i and j, respectively, while v± is the sum of all positive or all negative connection weights in the network.

To account for the near degeneracy of the modularity landscape (i.e., many near-optimal ways of partitioning a network into non-overlapping communities, Good et al., 2010) and for changes in community structure due to variations in the estimation parameters, the community detection algorithm was each initiated 100 times for three values of the spatial resolution parameter, centered around the default value of 1 (cf. Betzel & Bassett, 2017; Braun et al., 2015; Chen et al., 2016). Based on the results of these analyses, run separately for each of the three spatial resolution values, a consensus partition (i.e., whole-brain division into constituent communities) was estimated for each participant in each movie window (cf. Bassett et al., 2013; Lancichinetti & Fortunato, 2012).

##### Functional brain reorganization: Window-to-window versus context-based

Using the Network Community Toolbox, we estimated similarity in functional brain organization (i.e., community structure derived with the Louvain algorithm, as described above) between consecutive windows based on the adjusted normalized mutual information index [AMI], corrected for chance (Vinh, Epps, & Bailey, 2010). An index of window-to-window functional brain reorganization was computed by subtracting from 1 the average AMI across all pairs of temporally adjacent windows. This index combines spontaneous (i.e., stimulus-independent) window-to-window functional reconfiguration with reconfiguration driven by low-level, window-to-window perceptual fluctuations (e.g., presence/absence of objects, people).

Employing the event boundaries identified by independent raters in Ben-Yakov and Henson (2018) with a keypress when they felt that “one event [meaningful unit] ended and another began”, we selected pairs of non-overlapping 40 s windows, separated by ~ 5 to 7 s, which belonged to adjacent narrative segments (12 windows in total). Functional brain reorganization in response to high-level narrative context boundaries was estimated by subtracting from 1 the average AMI across all such pairs of temporally adjacent windows. This index reflects functional reconfiguration related to event boundaries, as well as stimulus-independent and lower-level reconfiguration indicative of featural changes (e.g., presence/absence of objects, people).

##### Network-based diversity in functional interactions

The participation coefficient assesses the diversity of a node’s intermodular connections (i.e., the extent to which a node interacts with nodes outside its native community) (Chen et al., 2016; Rubinov & Sporns, 2010). Here, a node’s native community was the one to which it was assigned in Power et al. (2011), the study that validated the functional atlas. It is worth pointing out that because our participation coefficient is based on a static community structure, it is not subject to the drawbacks associated with the participation coefficients derived from temporally varying communities (Thompson et al., 2020).

A node’s participation coefficient was based on the consensus partitions corresponding to each of the 177 sliding windows and was given by the formula

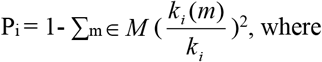

P_i_ is the participation coefficient of node i, M is the set of communities from a given partition (in our case, the whole-brain partition into communities, as described by Power et al. [2011]), k_i_(m) corresponds to the number of times that node i and all the nodes in community m have been assigned to the same community across all time windows, and k_i_ is the number of times that node i and the remaining 228 nodes have been assigned to the same community across all time windows. In our case, higher participation coefficients characterised nodes that tended to show an equal number of interactions (where interaction means assignment to the same community within a sliding window) with nodes from all the functional networks identified by Power et al.

### Brain-behavior analyses

#### Canonical correlation analysis (CCA)

To characterize the relationship between functional brain reorganization (i.e., changes in community structure) and network-level diversity in functional interactions, we used canonical correlation analysis (CCA, Hotelling, 1936) with cross-validation procedures (cf. Hair et al., 2008). CCA is a multivariate technique, which seeks maximal correlations between two sets of variables by creating linear combinations (i.e., canonical variates) from the variables within each set. Recently, CCA has been successfully used to investigate brain-behavior relationships in large datasets (see Smith et al., 2015; Tsvetanov et al., 2016; Wang et al. 2020). CCA was implemented in Matlab using the canoncorr module. In order to obtain reliable estimates of correlations between the brain or behavioral variables and their corresponding variates, it is generally recommended that CCA be performed on a sample size at least ten times the number of variables in the analysis (Hair, Anderson, Tatham, & Black, 1998), a criterion which was exceeded in all analyses reported below.

The performance of our CCA-derived models was tested by using a 10-fold cross validation procedure. Specifically, in the CCAs involving age, network participation and functional brain reconfiguration, the data were broken down into ten folds, all but two containing 64 participants for a total of 642 participants. For the CCAs involving fluid and crystallised intelligence, all but three of the ten folds of testing data contained 61 participants for a total of 613 participants. For both sets of CCAs, discovery analyses were conducted on nine folds of data and the resulting CCA weights were employed to derive predicted values of the brain and behavioral variate in the left-out (“test”) fold. This procedure was repeated until each of the ten folds served as “test” data once. The correlation between the predicted brain and behavioral variates across all testing folds was evaluated using a permutation test with 100,000 samples (cf. Smith et al., 2015).

To describe the relationship between the behavioral or brain variables and their corresponding variates across all the testing folds, we include correlations between the observed value of a brain or behavioral variable and the predicted value of its corresponding variate, as well as standardized coefficients, analogous to multiple regression coefficients, which indicate the unique association between the observed value of a behavioral or brain variable and the predicted value of its corresponding variate. For each of the two sets of CCAs, when evaluating the relationship between the predicted values of the two variates or the relationship between the actual value of each variable and the predicted value of its corresponding variate, we controlled for gender, handedness, subject-specific motion and subject-specific ROI functional homogeneity (RSC). 95% confidence intervals (CI) for each correlation and standardized regression-like coefficient were obtained by using the bootci function in Matlab (with default settings and 100,000 bootstrap samples).

The correlation and standardized regression-like coefficients described above are analogous to canonical loadings and canonical weights, respectively (see also Tsvetanov et al., 2016; Vatansever et al., 2017), with the only difference being that they are computed in the test, rather than the discovery, folds and, thus, reflect more conservative effect estimates.

#### Partial least squares analysis (PLS)

To identify patterns of ROI community participation that are specific to functional brain reorganization related to low-level featural versus high-level contextual changes/narrative event boundaries, we used partial least squares correlation often referred to as *PLS* (Krishnan et al., 2011), a multivariate technique that can identify in an unconstrained, data-driven manner, neural patterns (i.e., latent variables or LVs) related to individual differences variables (behavioral PLS). PLS was implemented using a series of Matlab scripts, which are available for download at https://www.rotman-baycrest.on.ca/index.php?section=345. In the behavioral PLS analyses we conducted, one matrix comprised residual scores on functional brain reorganization linked to event boundaries (i.e., average [1-AMI] across all neighbouring windows from different narrative segments from which age and average window-to-window functional brain reorganization were partialled out) (PLS 1) or average window-to-window functional brain reorganization (i.e., average [1-AMI] across all temporally adjacent windows from which age and average functional brain reorganization linked to event boundaries were partialled out) (PLS 2), whereas the second matrix contained each participant’s ROI participation matrices (Krishnan et al., 2011). Each matrix entry corresponded to the participation coefficient of one ROI from one subject.

In all the reported analyses, the significance of each LV was determined using a permutation test with 100000 permutations (in the permutation test, the rows of the ROI participation data are randomly reordered, Krishnan et al., 2011). In the case of our present analyses, PLS assigned to each ROI a weight, which reflected the respective ROI’s contribution to a specific LV. The reliability of each ROI’s contribution to a particular LV was tested by submitting all weights to a bootstrap estimation (100000 bootstraps) of the standard errors (SEs, Efron, 1981) (the bootstrap samples were obtained by sampling with replacement from the participants, Krishnan et al., 2011). We opted to use 100000 permutations and 100000 bootstrap samples (the same value used for the other bootstrapping and permutation-based testing herein reported) in order to increase the stability of the reported results, since these parameters are several orders greater than the standard ones (i.e., 500 permutations/100 bootstrap samples), recommended by McIntosh and Lobaugh (2004) for use in PLS analyses of neuroimaging data. A bootstrap ratio (BSR) (weight/SE) of at least 3 in absolute value was used as a threshold for identifying those ROIs that made a significant contribution to the identified LVs. The BSR is analogous to a z-score, so an absolute value greater than 2 is thought to make a reliable contribution to the LV (Krishnan et al., 2011), although for neuroimaging data BSR absolute values greater than 3 tend to be used (McIntosh & Lobaugh, 2004).

### Results

#### Window-to-window and narrative context-based brain reorganization increase with age; neither is linked to intelligence beyond age

We ran ten discovery CCAs to characterise the relationship of window-to-window and narrative-based functional reconfiguration with age, as well as fluid and crystallised intelligence. The discovery CCAs identified one significant mode, which was validated across all test sets (*r* of .15, *p* = 7 * 10^-5^). This mode indicated that greater functional brain reconfiguration (i.e., reduced similarity in community structure between temporally adjacent windows, as well as neighbouring windows from distinct narrative segments) typifies older individuals with lower fluid intelligence scores (see Figure 3-a, d). An inspection of the standardized coefficients revealed that the link between greater architectural variability and intelligence is mostly due to age (i.e., older individuals tend to have lower fluid intelligence scores relative to younger individuals, see Figure 3-b, e).

**Figure 3.**
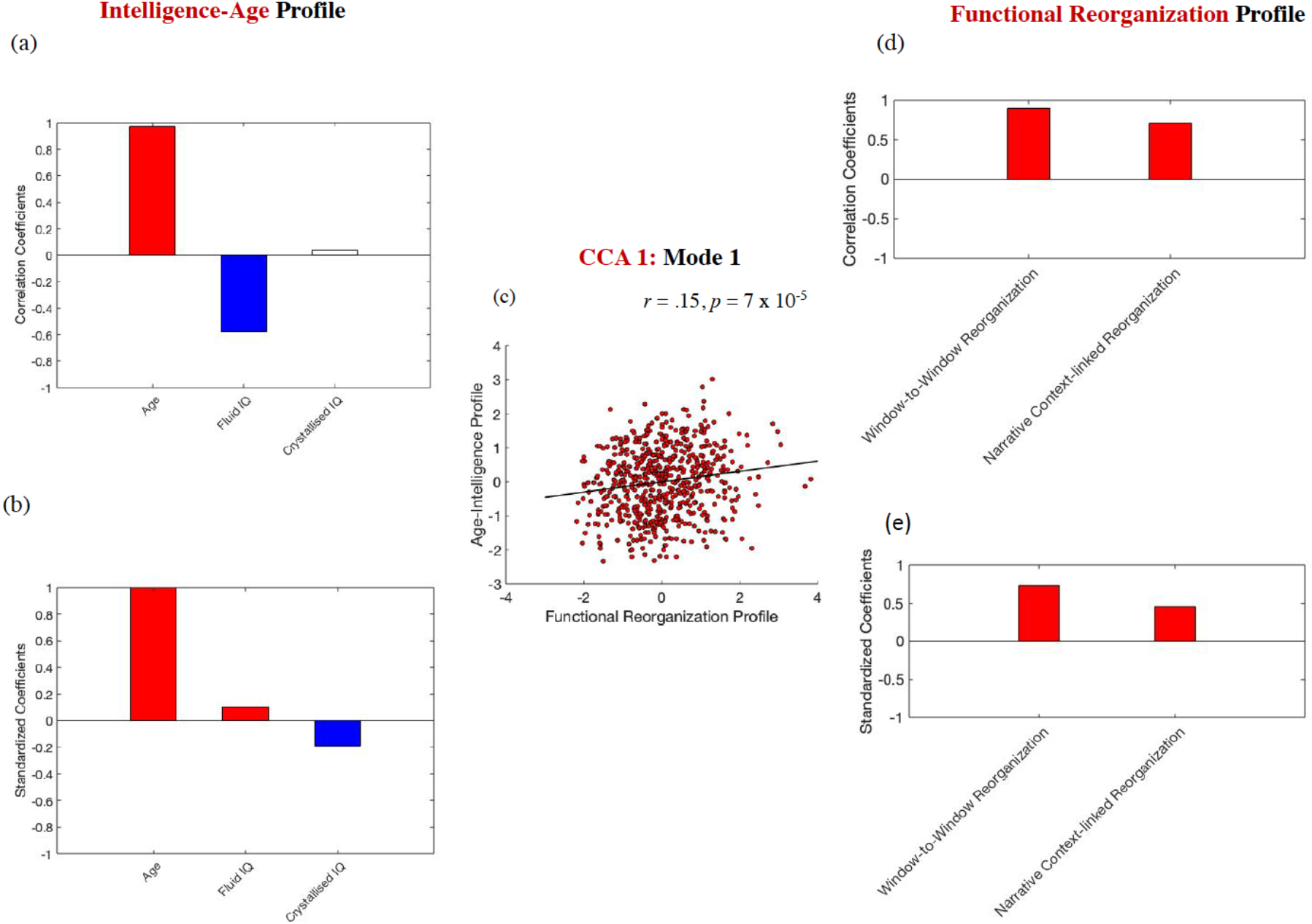
*Age, intelligence and functional brain reconfiguration.* The scatter plot in panel (c) is based on standardized variables. In panel (a), white boxes correspond to coefficients that were not robust across all the bootstrapping samples. Gender, handedness, ROI homogeneity and the summary motion metric were introduced as covariates.

#### Window-to-window and narrative context-based brain reorganization are linked to distinct patterns of network participation as a function of age

Ten discovery CCAs were conducted to probe the relationship between diversity in the functional interactions of the ten networks from the Power atlas (i.e., the average participation coefficient across all the ROIs within each network) and age-linked patterns of window-to-window versus narrative context-based brain reorganization. The discovery CCAs detected two significant modes, which were validated across all test sets (*r*s of .50 and .31, respectively, both *p*s of 10^-5^). The first mode indicated that, at older ages, functional brain reorganization is associated with greater participation of networks involved in self-guided cognition and creation of situation models during event perception (DMN), as well as top-down control (FPC) and attention (DAN), but reduced participation of networks implicated in environmentally driven processing (CON, AUD, SAL and VAN, see Figure 4). The second mode suggested that, at younger ages, stronger brain reconfiguration was linked to greater global participation, but particularly for the network involved in environmental vigilance and control maintenance (CON, cf. Figure 5).

**Figure 4.**
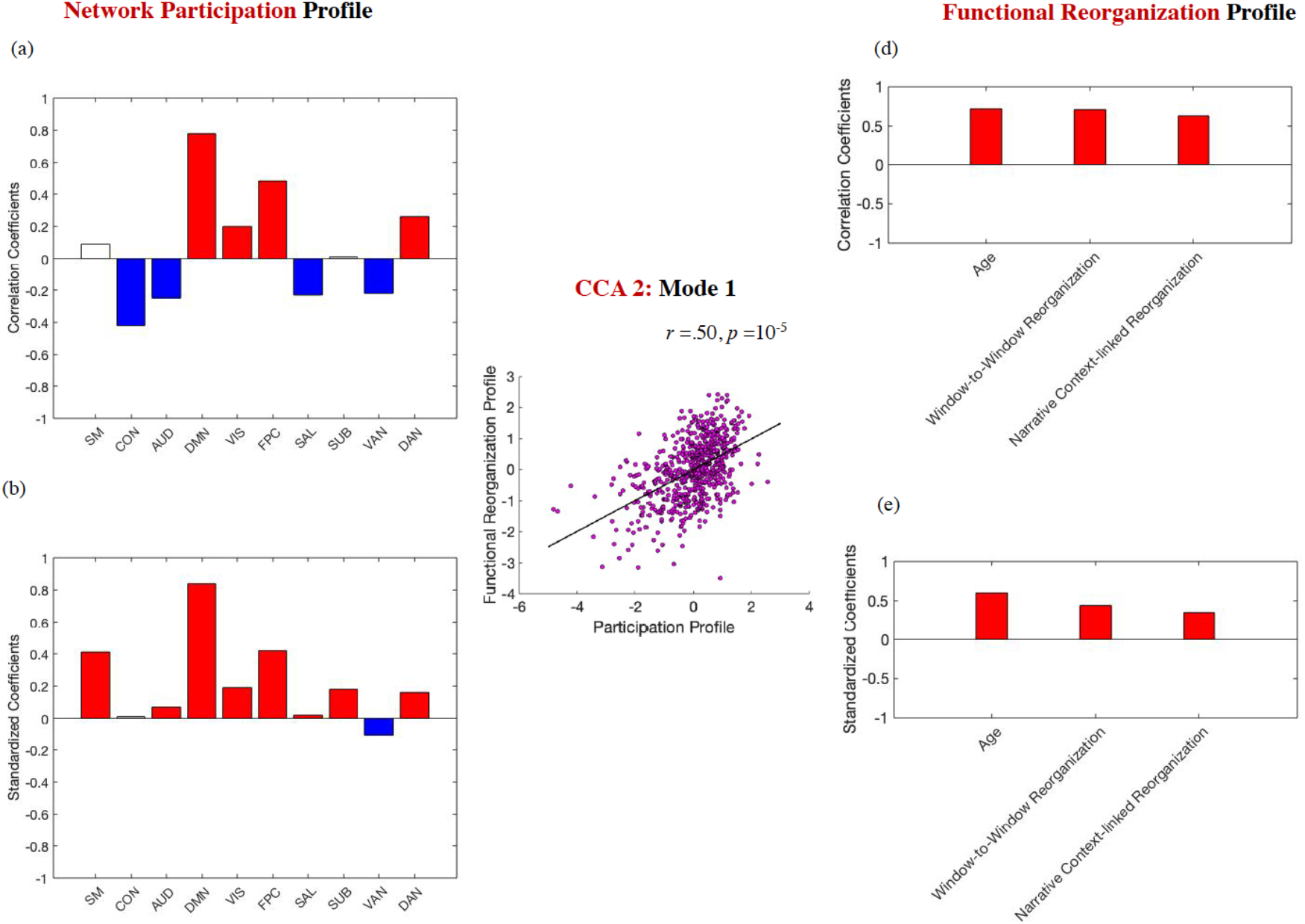
*Functional brain reconfiguration and network participation in older adulthood.* The scatter plot in panel (c) is based on standardized variables. In panels (a) and (b), white boxes correspond to coefficients that were not robust across all the bootstrapping samples. Gender, handedness, ROI homogeneity and the summary motion metric were introduced as covariates.

**Figure 5.**
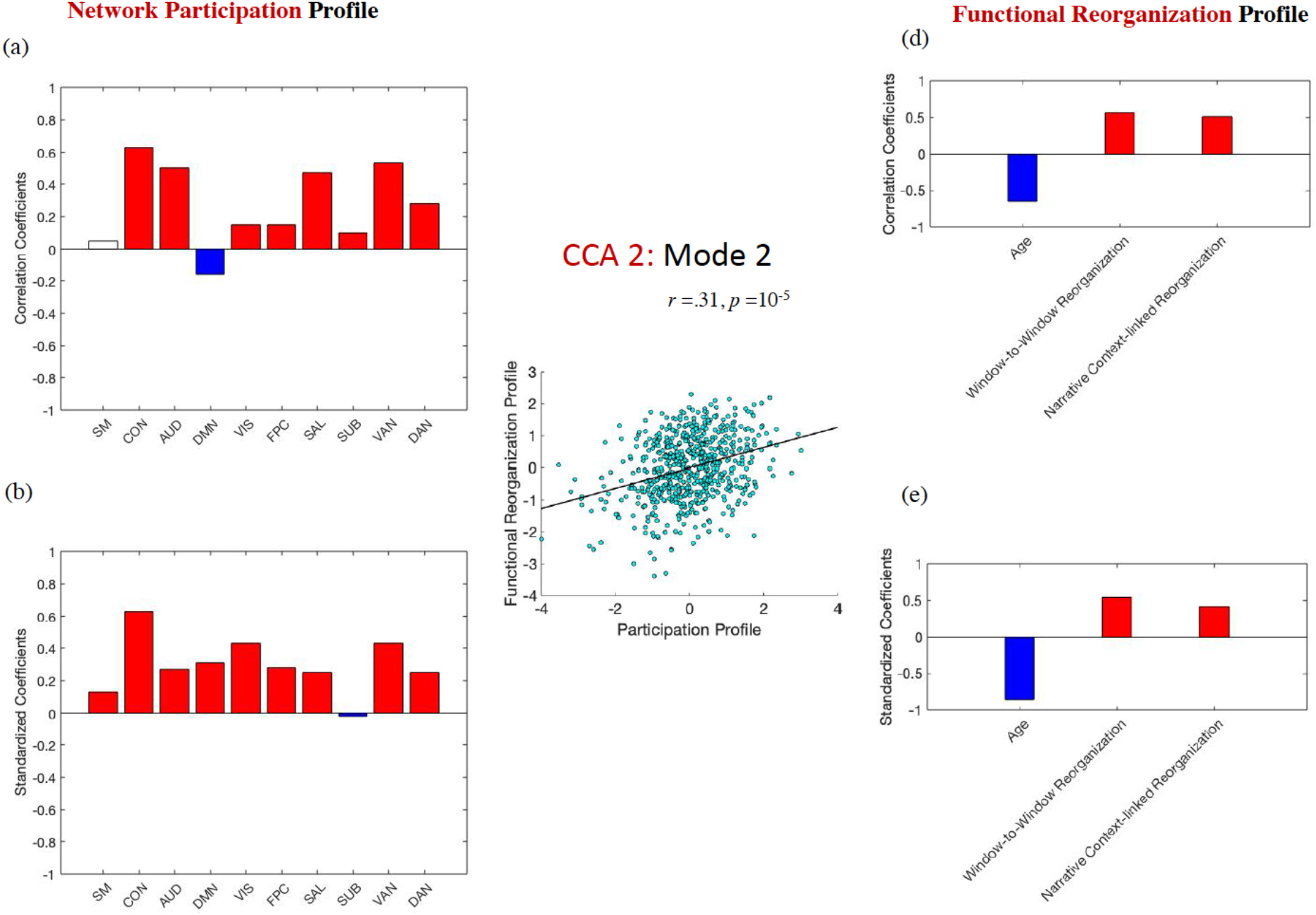
*Functional brain reconfiguration and network participation in younger adulthood.* The scatter plot in panel (c) is based on standardized variables. In panel (a), white boxes correspond to coefficients that were not robust across all the bootstrapping samples. Gender, handedness, ROI homogeneity and the summary motion metric were introduced as covariates.

#### The network participation profile linked to brain reconfiguration at younger ages, predicts fluid IQ independent of age and beyond levels of window-to-window /context - based reconfiguration; neither network participation profile predicts crystallised IQ

A robust regression analysis, conducted in Matlab with default settings (bisquare robust fitting weight function with a tuning constant of 4.685) and using fluid IQ as the outcome, revealed its significant positive association with the network participation profile linked to brain reconfiguration during, younger (*b* = .083, *SE* = .031, *t* (602) = 2.643, *p* =.008), but not older (*b* = −.053, *SE* = .037, *t*(602) = −1.433, *p* =.153), ages (covariates included window-to-window and context-based brain reconfiguration, respectively, age, sex, handedness, crystallised IQ, the summary motion metric and the summary ROI homogeneity metric) (see Figure 6). The corresponding robust regression analysis using crystallised IQ as the outcome unveiled no significant associations with either participation profile (both *p*s > .66). In neither robust regression analysis did levels of window-to-window or context-based functional reconfiguration made a significant contribution to either fluid or crystallised intelligence (all *p*s > .25).

**Figure 6.**
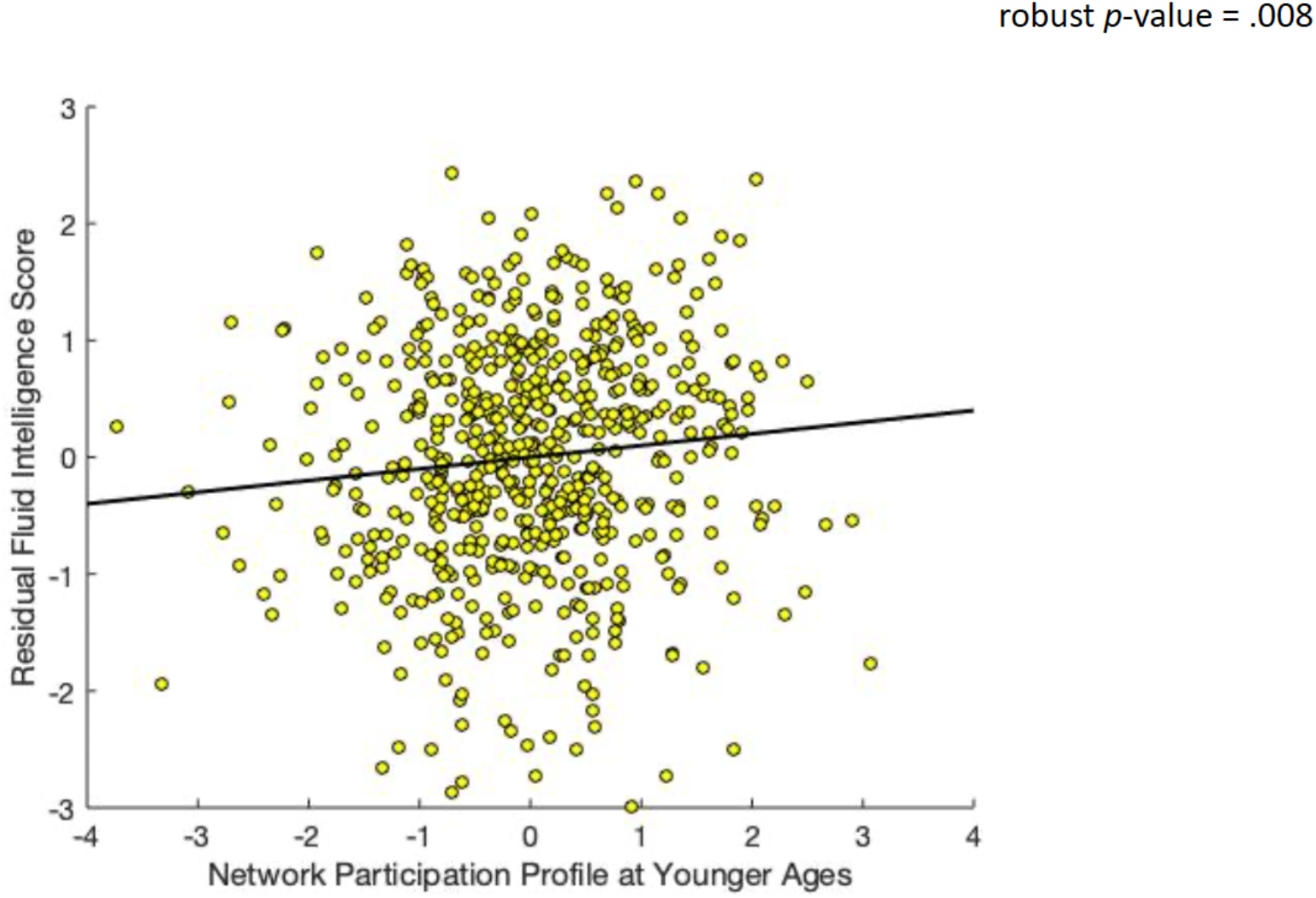
Scatterplot describing the association between the fluid intelligence score and the network participation profile linked to functional reconfiguration in younger adulthood, after controlling for all the covariates described in the text.

#### A subset of DMN ROIs predicts reconfiguration linked to narrative context boundaries, independent of age and window-to-window reconfiguration level

The behavioral PLS analysis identified one ROI participation LV (*p* = .0005) which was significantly linked to reorganization in response to movie event boundaries, independent of age and window-to-window brain reconfiguration levels (*r* = .25, 99% CI= [.25; .45]). Ten ROIs, all but one in the DMN, made a reliable contribution (absolute value BSR > 3) to this LV (insula, angular gyrus [AG], middle temporal gyrus [MTG], posterior cingulate cortex [PCC], superior frontal gyrus [SFG], dorsomedial and ventromedial prefrontal cortex [dmPFC, vmPFC], see Figure 7-a).

**Figure 7.**
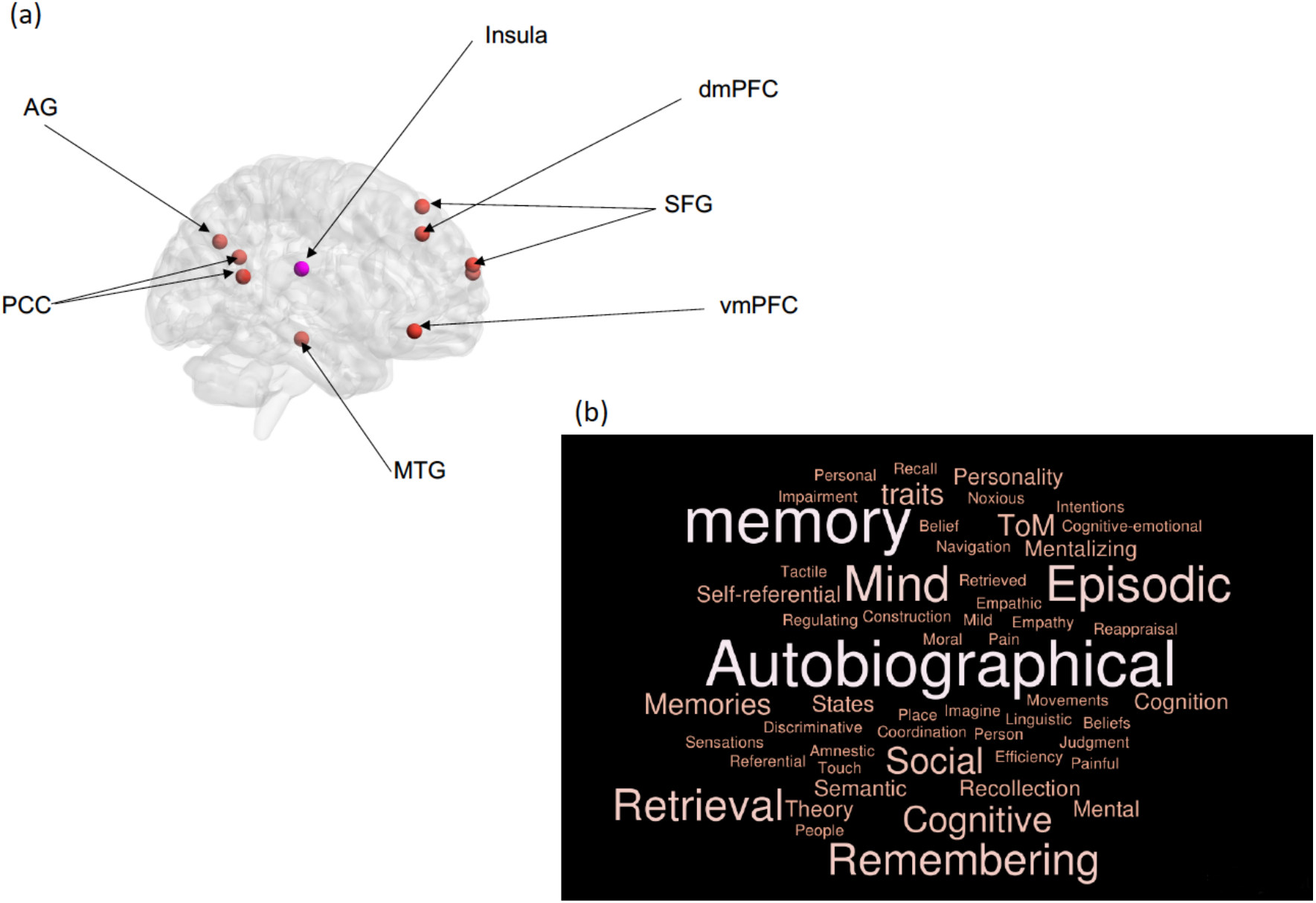
Panel (a). Results of the PLS analysis, showing the ROIs that demonstrated robust associations with context-based functional brain reconfiguration across the lifespan. The brain regions were visualized with the BrainNet Viewer (http://www.nitrc.org/projects/bnv/) (Xia et al., 2013). ROI colours reflect Power et al.’s network assignments (orange, DMN; pink, AUD). Panel (b) Results of the Neurosynth decoding analysis based on the ROIs in panel (a).

Subsequently, we conducted a decoding analysis in Neurosynth (Yarkoni et al., 2011), focused on the central voxel within each of the ROIs robustly linked by PLS to brain reconfiguration in response to event boundaries, in order to shed some light on their previously documented functional associations. As can be seen in Figure 7-b, the analysis revealed that the strongest z-score-based (Neurosynth z-scores > 4) associations were with “memory”, “autobiographical”, “episodic”, “retrieval”, “mind” and “remembering”. These decoding results are compatible with the interpretation that brain sensitivity to narrative context boundaries is uniquely associated with greater functional integration of ROIs that are relevant to the formation of ongoing event representations and play a key role in internally guided mnemonic processes (Honey, Newman, & Schapiro, 2017; Stawarczyk, Bezdek, & Zacks, 2019).

#### A subset of DMN, VIS and FPC ROIs predicts window-to-window reconfiguration, independent of age and event boundary-based reconfiguration level

This second behavioural PLS analysis identified one ROI participation LV (*p* = .0002) which was significantly linked to window-to-window reorganization, independent of age and narrative context-based brain reconfiguration levels (*r* = .26, 99% CI= [.26; .46]). Nine ROIs, encompassing the DMN (dmPFC, SFG), VIS (fusiform gyrus [FG], inferior occipital gyrus [IOG]), and FPC (inferior parietal lobule [IPL], medial frontal gyrus [MFG]) made a reliable contribution (absolute value BSR > 3) to this LV (see Figure 8-a).

**Figure 8.**
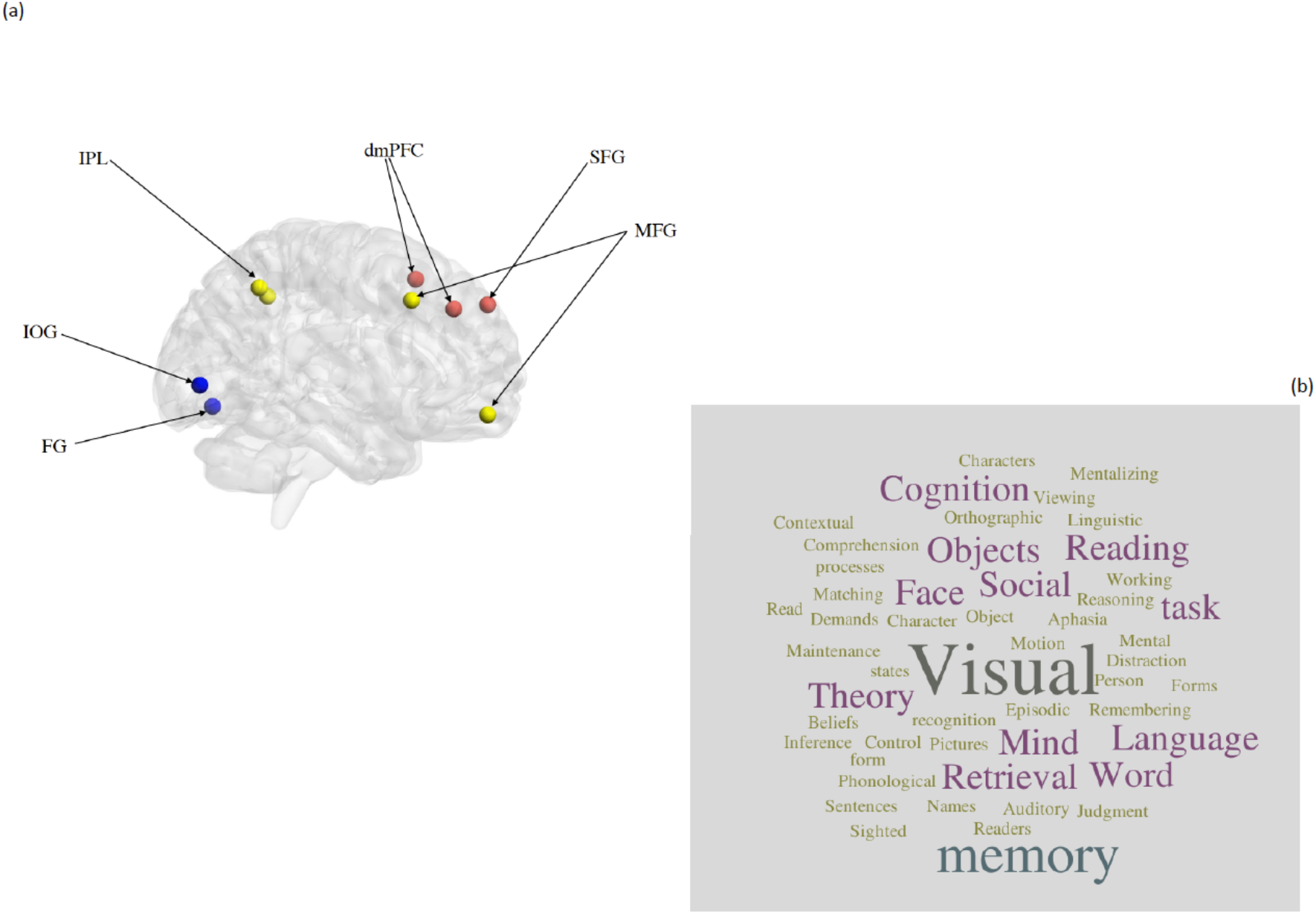
Panel (a). Results of the PLS analysis, showing the ROIs that demonstrated robust associations with window-to-window functional brain reconfiguration across the lifespan. The brain regions were visualized with the BrainNet Viewer (http://www.nitrc.org/projects/bnv/) (Xia et al., 2013). ROI colours reflect Power et al.’s network assignments (orange, DMN; blue, VIS; yellow, FPC). Panel (b) Results of the Neurosynth decoding analysis based on the ROIs in panel (a).

A decoding analysis in Neurosynth, similar to the one conducted for narrative context-based reconfiguration, revealed that the ROIs linked to window-to-window brain reconfiguration showed strongest functional associations with “visual”, “memory”, “objects”, “language”, “social”, “mind”, and “retrieval” (see Figure 8-b). This functional web was quite distinct from the one observed for the ROIs uniquely associated with context-based reconfiguration.

## Part 2: HCP Sample

### Participants

This sample included 176 unrelated participants, whose data had been released as part of the HCP 1200 subjects data package in March 2017. This sample represented the largest number of participants from the HCP 1200 subjects data release who were unrelated to one another and who had available data on all the demographic, behavioral and fMRI assessments of interest.

The majority of participants (N = 163) were right-handed. The sample included 70 younger men (21 between 22 and 25, 35 between 26 and 30, and 14 between 31 and 36 years of age) and 106 younger women (1 between 22 and 25, 49 between 26 and 30, and 56 between 31 and 36 years of age). Although age is presented here in the range format, as advocated by the HCP team (see Van Essen et al., 2012 for the rationale behind this age reporting strategy in HCP data releases), all our brain-behavior analyses used participants’ actual age in years, as available in the HCP restricted data release.

All participants were screened for a history of neurological and psychiatric conditions and use of psychotropic drugs, as well as for physical conditions or bodily implants that may render their participation unsafe. Diagnosis with a mental health disorder and structural abnormalities, as revealed by the MRI structural scans, were also exclusion criteria. Participants provided informed consent in accordance with the HCP research ethics board.

### Out-of-scanner measures

#### Fluid IQ

Form A of an abbreviated measure of the Raven’s Progressive Matrices (RPM; Bilker et al., 2012), a non-NIH Toolbox measure, gauged participants’ abstract reasoning skills (i.e., nonverbal fluid IQ). This task features patterns made up of 2×2, 3×3 or 1×5 arrangements of squares, with one of the squares missing. Participants must select one of five response choices that best fits the missing square on the pattern. The task has 24 items and 3 bonus items, arranged in order of increasing difficulty. The task is stopped though if the participant makes 5 consecutive incorrect responses. In line with existing guidelines (Bilker et al., 2012; Gray, Chabris, & Braver, 2003; Gray et al., 2005), total number of correct responses was used as a measure of abstract reasoning.

#### Subclinical depression and anxiety

To assess relatively stable subclinical variations in depression and anxiety, we used participants’ scores on the DSM-oriented depression and anxiety scales (see below for details). Both sets of scores were derived from participants’ responses to relevant items on the Achenbach Adult Self-Report (ASR) instrument for ages 18–59 (Achenbach, 2009). The ASR contains a total of 123 statements relevant to psychological functioning and requires participants to rate on a 3-point scale (0 *not true,* 1 somewhat or sometimes true, 2 *very true or often true)* how well each item described them over the previous six months. The DSM-oriented depression scale includes items such as “I am unhappy, sad, or depressed”. The DSM-oriented anxiety scale includes items such as “ I worry about my future”.

### Out-of-scanner control measures

#### Current negative emotion experience

Participants completed the NIH Toolbox Negative Affect Survey, which assesses separately current levels of experienced sadness (e.g., “I felt sad.”, “I felt like a failure.”), anger (e.g., “I felt angry.”, “I felt bitter about things.”), and fear (e.g., “I felt frightened.’, “I had a racing or pounding heart.”), respectively. The measure requires participants to rate on a 5-point scale (1 *never* to 5 *always)* how often they experienced the relevant emotion within the past seven days. Scores on the sadness, anger and fear subscales were averaged to create an index of current negative emotional state, which was entered as a covariate in all the hypothesis testing analyses.

### In-scanner task

Participants completed four movie viewing runs over two scanning sessions on two separate days. Each run was about 15 minutes long and comprised 1 to 4.3 minutes long excerpts from Hollywood movies, as prepared and published by Cutting, Brunick, and Candan (2012), and independent films, freely available under Creative Commons Licensing. All four movie runs include a Vimeo repeat clip at the end, which was not included in the analyses. In each run, 20 s of rest (i.e., black screen with “REST” in white text) precede the beginning and follow the end of each movie clip.

#### Movie features

To quantify low-level featural fluctuations in the movie task, we used the output of the semantic feature coding conducted by Jack Gallant’s laboratory (cf. Huth, Nishimoto, Vu, & Gallant, 2012) and available as part of the HCP1200 Subjects Data Release. By using this semantic feature coding, we were able to characterise the extent to which window-to-window reconfiguration in functional architecture is yoked to ongoing, meaningful fluctuations in the external environment. Each movie frame is coded for the presence/absence (0/1) of 859 semantic elements recorded as nouns (N = 629), verbs (N = 229) and adjective(s) (N =1). Our analyses focused on semantic features listed as nouns, which we took to reflect environmental attributes (e.g., presence of objects, people, buildings) and semantic features listed as verbs, which reflected actions performed by the movie characters or natural phenomena. As we detail below, the brain data corresponding to each movie was broken down into 40 s second sliding windows, moved in increments of 2 s. The binary semantic feature matrices corresponding to each sliding window from the brain data were averaged. In these averaged matrices, the value in each cell signified the percentage of frames within a sliding window when an entitity or action was present. As in the brain data, window-to-window similarity in semantic features was indexed with the adjusted mutual information index (AMI) (Vinh et al., 2012), with each unique value in the averaged semantic features for each window acting as a community label.

### fMRI data acquisition

Images were acquired with a customized Siemens 3T “Connectome Skyra” scanner housed at Washington University in St. Louis (32-channel coil). Pulse and respiration were measured during scanning. Functional images were acquired with a multiband EPI sequence (TR=1000 ms, TE=22.2 ms, flip angle=45°, FOV = 208 x 208 mm, 85 slices of 1.6 × 1.6 mm in-plane resolution, 1.6 mm thick, no gap). Two of the movie runs were acquired with an anterior-to-posterior, while the other two with a posterior-to-anterior, phase encoding sequence (so that phase encoding sequence effects could cancel each other out over the all runs).

### fMRI data preprocessing

The present report used the minimally preprocessed movie watching data from the HCP 1200 subjects data release. These data have been preprocessed with version 3 of the HCP spatial and temporal pipelines (Smith et al., 2013; for specification of preprocessing pipeline version, see http://www.humanconnectome.org/data). Spatial preprocessing involved removal of spatial and gradient distortions, correction for participant movement, bias field removal, spatial normalization to the standard Montreal Neurological Institute (MNI)-152 template (2 mm isotropic voxels), intensity normalization to a global mean and masking out of non-brain voxels. Subsequent temporal preprocessing steps involved weak high-pass temporal filtering with the goal of removing linear trends in the data.

#### Additional denoising

Because motion can significantly impact functional connectivity measures (Power et al., 2012; Van Dijk et al., 2012), we implemented the same additional preprocessing steps used to address this potential confound in the Cam-Can data. Specifically, in the CONN toolbox (version 17c; Whitfield-Gabrieli & Nieto-Castanon, 2012), linear regression was used to remove from the BOLD time series of each ROI the BOLD time series of the voxels within the MNI-152 white matter and CSF masks, respectively (i.e., the default CONN option of five CompCor-extracted principal components for each, Behzadi, Restom, Liau, & Liu, 2007), the 6 realignment parameters, their first-order temporal derivatives and their associated quadratic terms (24 regressors in total, cf. Bolt et al., 2017), as well as the main movie effects, obtained by convolving a boxcar task design function with the hemodynamic response function, and their first temporal order derivative (cf. Braun et al., 2015; Vatansever et al., 2015; Westphal, Wang, & Rissman, 2017). The last denoising step was implemented in order to isolate movie-related functional coupling from mere co-activation effects corresponding to the beginning and end of a movie clip (i.e., two regions that are both activated at the beginning of a movie clip and de-activated at its end, although they do not “communicate” with one another throughout the movie clip). The residual BOLD time series were bandpass filtered (0.008 Hz< f < 0.09 Hz), linearly detrended and despiked (all three are default CONN denoising steps). Following these corrections (which again did not include global signal regression), an inspection of each subject’s histogram of voxel-to-voxel connectivity values revealed a normal distribution, approximately centered around zero, which would suggest reduced contamination from physiological and motion-related confounds (cf. Whietfield-Gabrieli & Castanon, 2012). Nonetheless, in supplementary analyses, accompanying all the brain-behavior tests, we confirmed that all the reported effects were not driven by individual differences in motion, as they remained unchanged after controlling for the average relative (i.e., volume-to-volume) displacement per participant, a widely used motion metric (Power et al., 2012, 2015; Satterthwaite et al., 2013).

### fMRI data analysis

#### ROI time series

We followed the same steps as in the Cam-Can data in order to extract the timeseries from 229 ROIs from the Power et al. (2011) atlas.

#### Functional connectivity analyses

Pairwise coupling among the 229 ROIs was estimated in CONN, separately for each of the 14 movies. Periods of rest between consecutive movie clips were eliminated from the analyses. As in the Cam-Can data, the pairwise correlations among all the ROIs were expressed as Fisher’s z-scores, all of which (i.e., positive and negative alike) were subsequently used to compute the indices of interest for all connectivity analyses.

To characterize individual differences in dynamic network structure, we broke down each movie clip into partially overlapping 40 s long windows for a total of 1152 windows. Similar to the Cam-Can analyses, pairwise coupling among the 229 ROIs was estimated in CONN using a sliding window of 40 s in length (40 volumes) with a two-TR gap in-between windows and a “hanning weighting” (i.e., greater weight to the scans in the middle of the window relative to the ones at the periphery) applied to all the time points within a window.

#### Network-level analyses

We followed the same steps as in the Cam-Can data with the following exceptions. We only computed similarity in functional brain organization (i.e., community structure) between consecutive windows, since, due to the shorter duration and the structure of the HCP movie clips, narrative event boundaries were less legible and, as such, did not constitute a point of inquiry. Instead, to index brain sensitivity to object-versus action-related changes, we computed the Spearman’s rank correlation between window-to-window similarity in functional brain organisation (i.e., community structure) and window-to-window noun-versus verb-based semantic similarity. Because we were specifically interested in brain- (noun/verb-based) movie *coupling*, overall window-to-window brain reconfiguration was regressed out from both indices. The two residual brain-movie (noun-vs. verb-based) couplings were used in all the reported analyses. Participation coefficients for each of the 229 ROIs were computed as in the Cam-Can dataset.

##### Reliability analyses

To test whether a unitary construct can be extracted for each neural index of interest across all 14 movies, we conducted separate reliability analyses on the 42 values associated with each index (i.e., three values for each of the 14 movies, corresponding to the community detection estimates obtained with a spatial resolution parameter of .95, 1, 1.05). Since subject motion can impact such reliability estimates, we present the relevant Cronbach’s alpha values, both before and after regressing out subject level average frame-to-frame displacement (see Preprocessing above for the additional motion effect removal procedures already implemented). Additionally, to reflect the variables used in our analyses, for the brain- (noun/verb-based) movie coupling, we present reliability estimates based on data from which we regressed out both subject motion and spontaneous window-to-window functional brain reconfiguration.

For the window-to-window brain organisation similarity index, we obtained Cronbach’s alphas of .87 and of .89 (with regression of the motion summary statistic). The brain-(noun) movie coupling index showed Cronbach’s alphas of .66 and of .68 (with regression of motion and window-to-window brain reconfiguration). For the brain-(verb) movie coupling index, we observed Cronbach’s alphas of .65 and of .62 (with regression of motion and window-to-window brain reconfiguration). Across all 229 ROIs, the participation coefficients showed Cronbach’s alphas between .76 and .98 (both with and without regression of the summary motion statistic).

### Brain-behavior analyses

#### Canonical correlation analysis (CCA)

We used canonical correlation analysis (CCA, Hotelling, 1936) with cross-validation procedures (cf. Hair et al., 2008) in order to characterize the relationships between our neural indices of interest (coupling between movie features and functional brain reorganization; ROI-specific diversity in functional interactions), as well as between the relevant brain (coupling between window-to-window changes in movie features and functional brain reorganization [dissimilarity in community structure]) and behavioral (fluid IQ, subclinical depression, subclinical anxiety) variables. Age was introduced in the CCAs in order to probe potential developmentally specific effects (Ofen et al., 2012; Petrican & Grady, 2017). As recommended in the literature (Hair et al., 1998), all CCAs were based on sample sizes more than ten times the number of variables in the analysis (Hair, Anderson, Tatham, & Black, 1998).

As in the Cam-Can data, the performance of our CCA-derived models was tested by using a 10-fold cross validation procedure. Specifically, the data were broken down into ten folds, six of which contained 18 participants (the remaining folds contained 17 participants each). The cross-validation procedure and relevant model descriptors were identical to those in the Cam-Can data. In both sets of CCAs, when evaluating the relationship between the predicted values of the two variates or the relationship between the actual value of each variable and the predicted value of its corresponding variate, we controlled for gender, handedness, subject-specific motion, years of education. In CCA 1 (see Figure 9) we additionally controlled for current negative emotional experience to ensure that the observed associations are not due to global negative mood around the time of testing.

**Figure 9.**
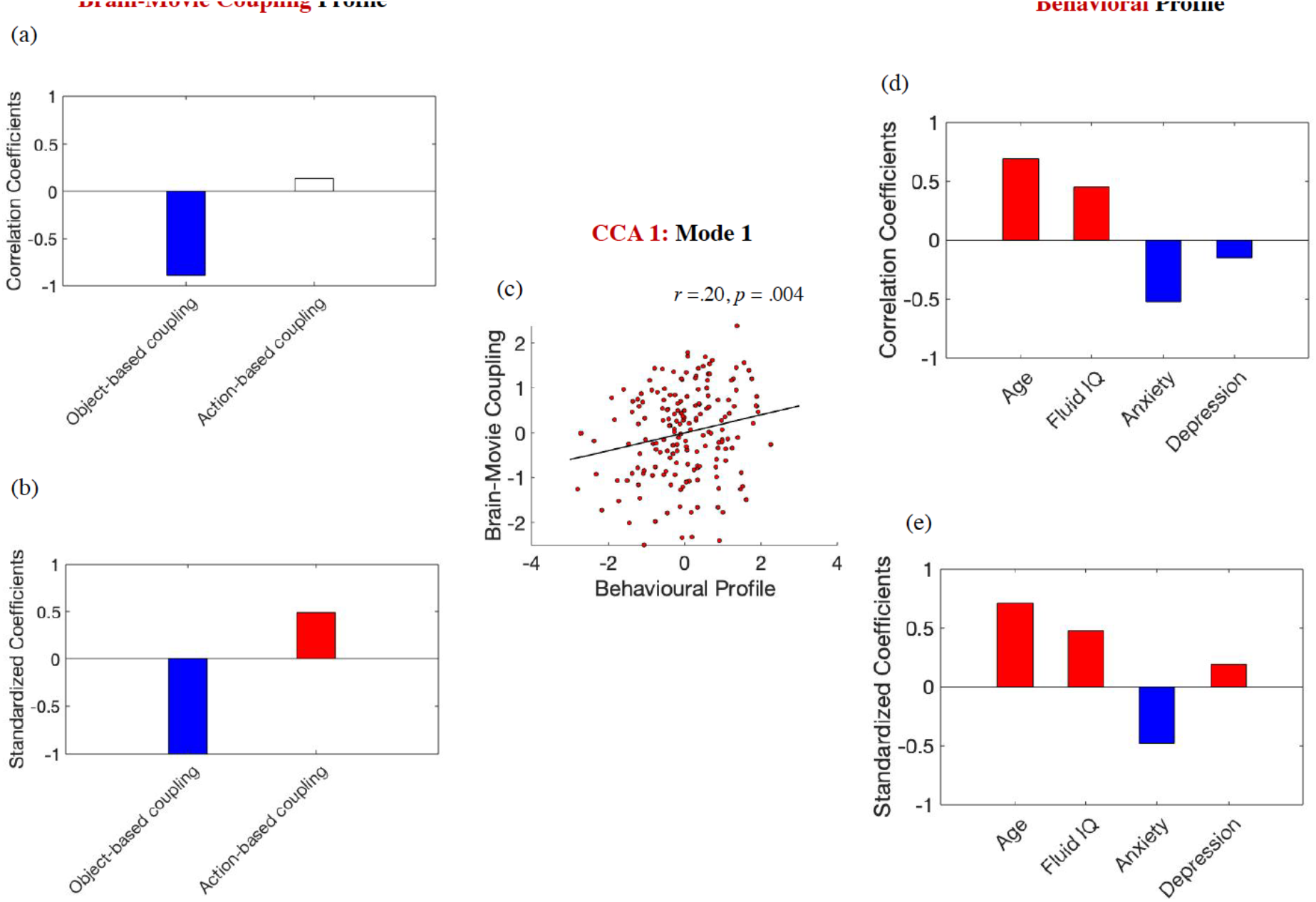
Brain-movie coupling and cognitive-affective function in the HCP sample. The scatter plot in panel (c) is based on standardized variables. In panel (a), white boxes correspond to coefficients that were not robust across all the bootstrapping samples. Gender, handedness, years of education, the summary motion metric and current negative emotional experience were introduced as covariates.

### Results

#### Anxious younger adults with lower fluid IQ show greater brain sensitivity to object-based, but not action-based changes in the external environment

Ten discovery CCAs were conducted to probe the relationship between age, fluid IQ, anxiety and depression, on one hand, and coupling of functional brain reconfiguration with semantic feature (noun vs. verb-based) changes across all 14 movies. The discovery CCAs detected one significant mode, which was validated across all test sets (*r* of .20, *p* = .004). This mode indicated that greater functional brain reconfiguration as a function of object-based, but not action-based changes, typifies anxious younger adults with lower fluid IQ scores (see Figure 9 for the correlation [panels a, d] and the standardized coefficients [panels b, e] of the ROI participation variables on their corresponding canonical variate across all test CCAs, as well as the scatter plot describing the linear association between the ROI participation variate and the brain-object-based movie coupling across all the “test” folds [panel c]).

#### Greater brain sensitivity to object-based environmental changes is linked to greater whole-brain participation of parietal and medial temporal areas involved in autobiographical and episodic memory

We conducted ten discovery CCAs, probing the link between brain-environment coupling based on object-based variations and participation of the eleven ROIs uniquely linked in the Cam-Can to brain reconfiguration evoked by high-level, narrative context changes. Because we were interested specifically in brain-environment coupling with respect to object-related variations (due to its relevance to cognitive-affective adaptation, as shown in the previous set of CCAs), we regressed out from the brain-index not only global window-to-window brain reconfiguration (as in the prior analysis), but also brain-environment coupling based on narrative action changes. Age was introduced in this analysis to probe whether its link to brain-environment coupling is independent of the narrative ROI participation patterns (in the Cam-Can, the ROI participation profiles were shown to contribute to high-level context-based reconfiguration irrespective of age, but it was unclear whether the same would be true with to lower-level featural fluctuations).

One significant mode emerged from the discovery CCAs, which was replicated across all test sets (*r* of .19, *p* = .005). This mode indicated that stronger brain-environment coupling (with respect to object-based fluctuations) was associated with greater participation across most of the Cam-Can ROIs at younger ages, but particularly the medial temporal and parietal ROIs, which, based on Neurosynth decoding, were most relevant to “memory”, “autographical”, “episodic” and “retrieval” (see Figure 10 for the correlation [panel a] and the standardized coefficients [panel b] of the ROI participation variables on their corresponding canonical variate across all test CCAs, the scatter plot describing the linear association between the ROI participation variate and the brain-object-based movie coupling across all the “test” folds [panel c], as well as the results of the Neurosynth decoding analysis constrained to the ROIs that showed robust correlations with the extracted covariate in panel [a]).

**Figure 10.**
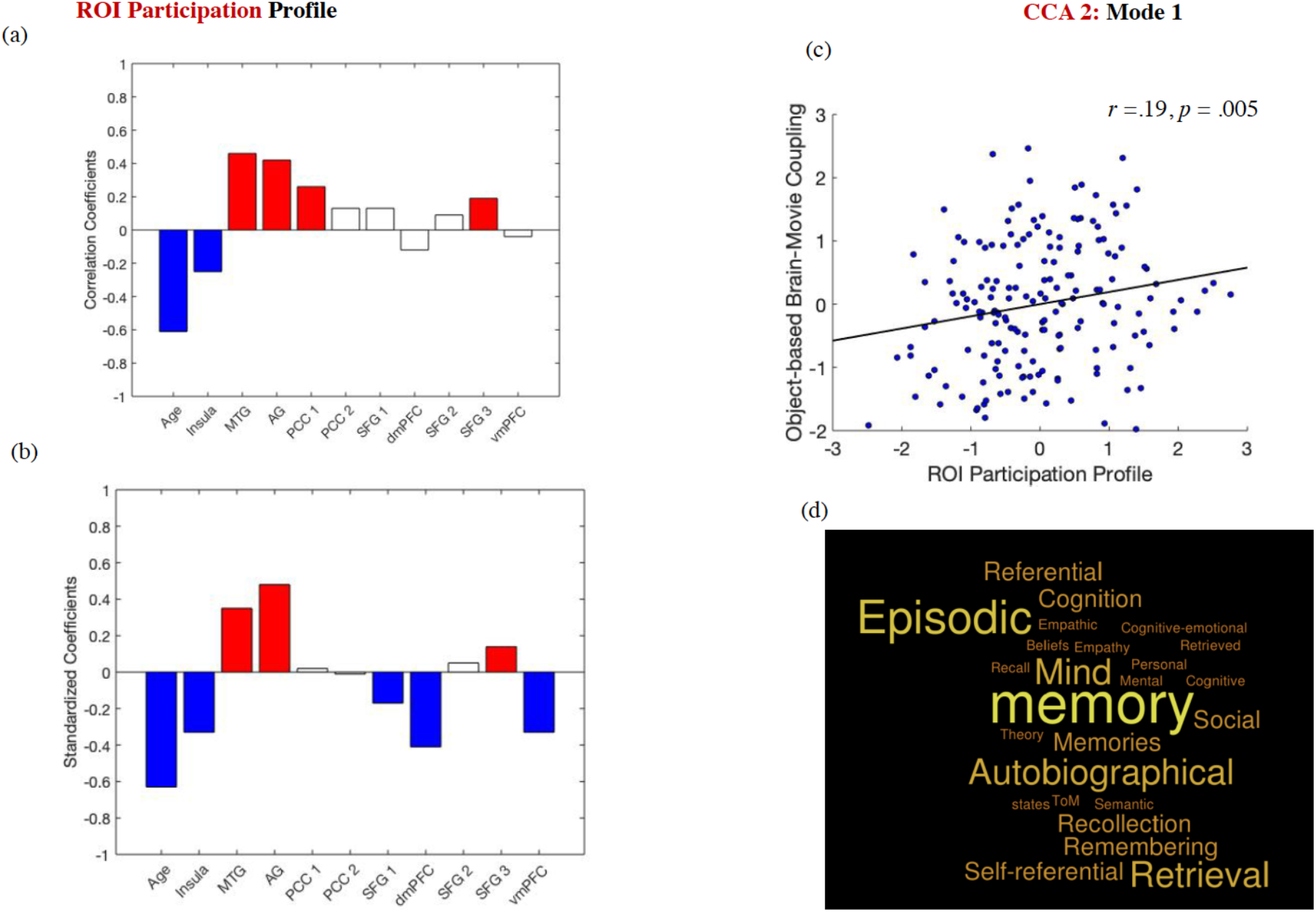
*The ROI participation profile linked to brain-object-based movie coupling.* The scatter plot in panel (c) is based on standardized variables. In panels (a) and (b), white boxes correspond to coefficients that were not robust across all the bootstrapping samples. Gender, handedness, years of education and the summary motion metric were introduced as covariates. Panel (d) presents the results of the Neurosynth decoding analysis based on the ROIs in panel (a).

## Discussion

Extending prior literature on the role of brain signal variability in fostering a more differentiated and flexible response to the environment (e.g., Garrett et al., 2013a, b, 2015, 2020; Grady & Garrett, 2018), we provide novel evidence that the adaptiveness of both moment-to-moment and context-based fluctuations in whole-brain functional architecture varies across the lifespan as a function of the underlying network communication profiles. We further demonstrate that enhanced brain-environment alignment with respect to concrete featural fluctuations is stronger at younger ages, but associated with adverse implications for both cognitive and affective adaptation (i.e., fluid intelligence, depression/anxiety). Finally, we document the network integration profiles that link functional brain reconfiguration at multiple timescales and are, thus, likely to be key to understanding the dynamics behind typical and atypical variations in event processing.

Using a naturalistic, dynamic cognition paradigm in an adult lifespan sample, we demonstrate that the adaptiveness of functional brain reconfiguration hinges on the associated patterns of whole-brain network participation. Specifically, superior cognitive adaptability (i.e., higher fluid intelligence, Cattell, 1971) was not linked to variability in functional neural architecture per se, but rather to a network participation profile implicated in dynamic brain reconfiguration in younger adulthood. This profile reflects a pattern of whole-brain functional integration anchored in networks implicated in vigilance and control maintenance (CON), as well as environmentally driven attention (VAN) and behavioural regulation (SAL) (see Figure 5-a, Corbetta & Shulman, 2002; Sadaghiani & D’Esposito, 2015; Seeley et al., 2007; Sridharan, Levitin, & Menon, 2008). These results thus dovetail nicely with findings from the literature on brain signal variability, thereby implying that the beneficial consequences of both signal and architectural variability are likely to involve neural tuning to the dynamics of the external world (Garrett et al., 2013a, 2020; Grady & Garrett, 2018). Our findings are also compatible with extant theory which suggests that flexibility in the interactions of cognitive control networks, particularly CON, are a key contributor to fluid intelligence (Barbey, 2018; Duncan et al., 2020). Overall, our results imply that adaptive patterns of dynamic brain reconfiguration during movie watching are linked to greater whole-brain informational flow through a system implicated in maintaining current task demands (i.e., CON, Dosenbach et al., 2006, 2008) and which is thus very well-positioned to support the “working event model” of an ongoing experience (Radvansky & Zacks, 2017).

Complementarily, the participation profile linked to variability in functional architecture during older adulthood was typified by reduced CON, VAN, and SAL participation and, instead, reflected most strongly the diverse interactions of the networks involved in self-guided cognition (DMN) and top-down control (FPC), in particular, but also of those implicated in goal-directed attention (DAN) and visual processing (Andrews-Hanna, Saxe & Yarkoni, 2014; Corbetta et al., 2000; Spreng et al., 2010) (see Figure 4-a). Our finding that functional brain reconfiguration in older adulthood depends on neural communication pathways grounded in the DMN and FPC complements current theories of cognitive ageing, which posit that age-related declines in the ability to engage strategically with the external environment in the here-and-now are compensated by drawing on accumulated world knowledge (Spreng & Turner, 2019). This age-related semanticisation, stemming from progressively stronger functional coupling between the DMN and the FPC (Turner & Spreng, 2015), helps preserve task performance in contexts where prior knowledge is relevant (Umanath & Marsh, 2014). One such context may be event segmentation, where performance preservation with ageing may be due to increasing reliance on semantic knowledge (rather than perceptual representations) during event perception (Radvansky & Dijkstra 2007). This conjecture is compatible with recent findings of ageing-related activity reductions in canonical episodic memory areas, but not schematic processing areas, in response to narrative event boundaries observed in the Cam-Can sample, an effect that emerges despite the lack of age-related differences in behavioural event segmentation (Reagh, Delarazan, Garber, & Ranganath, 2020). Importantly, though, we did not find a significant association between crystallized intelligence (i.e., semantic knowledge) and the participation profile typifying architectural variability in older age. Hence, rather than reflecting ageing-related semanticisation, this network profile may simply indicate ageing-related deficits in DMN disengagement, which may underpin its greater functional integration, as observed here (Samu et al., 2017). Further research is needed to shed light on the role of DMN during naturalistic cognition in older adulthood.

Complementing findings on the adaptiveness of modulating brain signal variability based on environmental complexity (e.g., Garrett et al., 2020), our study provided novel insights into the adverse functional implications linked to coupling in the dynamics of whole-brain architecture and concrete environmental features. Specifically, prior evidence indicates that anxiety and depression are associated with divergent processing biases, which impact event perception (Belzung et al., 2015; Brewin et al., 2010; Petrican et al., 2015; Sherrill et al., 2019). Accordingly, we find that subclinical anxiety is linked to increased brain-environment alignment, implying greater tuning into the dynamics of the external perceptual world (Figure 9-a, b), while an opposite tendency, consistent with attentional disengagement from concrete environmental dynamics in the here-and-now, is observed for subclinical depression (Figure 9-e). Brain sensitivity to concrete featural fluctuations was also associated with poorer fluid intelligence. This implies that environmental entrainment, a plausible marker of preferential reliance on sensory-bound, rather than more abstract mental representations, which is linked to high levels of both state and trait anxiety, may prevent successful strategic processing (Hermans et al., 2014; Matthews, Yiend, & Lawrence, 2006; Sylverster et al., 2012). Importantly, these effects were specific to brain sensitivity to object/spatial layout fluctuations and did not extend to brain sensitivity to action-based changes, which have been found to support typical event segmentation processes from childhood onwards (Levine, Buchsbaum, Hirsh-Pasek, & Golinkoff, 2019; Magliano & Zacks, 2011; Swallow, Kemp, & Simsek, 2018).

Our study also contributed novel evidence on the overlapping network communication profiles underlying brain-environment alignment with respect to both higher-level contextual and lower-level featural fluctuations. In both cases, brain-environment coupling was associated with greater whole-brain informational flow (i.e., participation) in a subset of canonical DMN ROIs, including the left AG, left MTG, PCC and left SFG, the majority of which had been implicated in event memory reactivation (Chen et al., 2017). The AG and PCC play key roles in recollection (Ranganath & Ritchey, 2012; Ritchey & Cooper, 2020; Rugg & Vilberg, 2013) and have been widely implicated in the integration of multimodal information at longer timescales, thereby supporting the creation of the so-called “situation/event models”(Bonnici, Richter, Yazar, & Simmons, 2016; Chen et al., 2016; Radvansky & Zacks, 2017; Stawarcyzk et al., 2019; Yazar, Bergstrom, & Simmons, 2017). The left AG, in particular, plays a causal role in episodic context creation during perception by acting as an online buffer for combining past and currently presented information (Branzi, Pobric, Jung, & Lambon Ralph, 2019; Humphreys & Lambon Ralph, 2015). Expanding this literature, we document the role of AG in integrating information across the whole brain during event perception in order to align the external environmental and internal neural dynamics. The AG seems thus to be well-placed to contribute to cognitive adaptability (i.e., fluid IQ), together with “multiple demand” cognitive control areas (Barbey, 2018; Duncan et al., 2020) by shifting attentional focus from a featural, sensory-bound to a more abstract processing mode in a context-specific manner.

The left MTG demonstrated a similarly robust association with functional brain reconfiguration triggered by both high and low-level environmental changes. Like AG, this region underpins updating of semantic features related to the present context (Branzi, Humphreys, Hoffman, & Lambon Ralph, 2020), while uniquely partaking into the controlled retrieval of semantic information (Hoffman, McClelland, & Lambon-Ralph, 2018). The MTG and AG could, plausibly, make complementary contributions to context creation and updating (Kurby & Zacks, 2008). Specifically, the AG-centered participation profile may provide the low-level, episodic perceptual detail from which a strong sense of sensory vividness and grounding in the here-and-now stem (Ramanan, Piguet, & Irish, 2018). Complementarily, the MTG-linked participation profile may support continual updating of the underlying semantic structure based on the influx of environmental information and controlled retrieval of already stored world knowledge, thereby synching the “working event model” with the ongoing experience (Hoffmann et al., 2018; Zacks, 2020).

Future studies are warranted to address limitations of our present research. First, use of a larger battery of movies (as in HCP), covering diverse artistic interests and production dates, with a strong narrative plot that allows reliable extraction of narrative event boundaries (as in Cam-Can), is needed to characterise hierarchical event perception dynamics. Second, inclusion of a full lifespan sample, as well as complementary cross-sectional/longitudinal designs may elucidate the role of brain-environment entrainment during developmental stages characterised by distinct learning needs (Baldwin & Kosie, 2020). Third, although behavioural event segmentation is largely preserved in healthy ageing (Kurby & Zacks, 2018; Reagh et al., 2020; Sargent et al., 2013), future studies probing the link between event segmentation performance and its neural substrates across the lifespan would be critical in furthering our understanding of developmental differences in information processing. Fourth, research employing alternate methods for estimating dynamic brain reconfiguration would elucidate the boundary conditions of the effects herein documented (Gonzalez-Castillo & Bandettini, 2018; Iraji et al., 2020).

In sum, we demonstrate that the adaptiveness of dynamic brain reconfiguration during naturalistic event cognition varies across the lifespan based on the associated patterns of network interaction. Specifically, similar to brain signal variability, the adaptiveness of architectural variability seems to hinge on its relevance to enhanced processing of the external world. Complementing the literature on environmental modulation of brain signal variability, we further show that brain-environment alignment with respect to concrete, featural fluctuations is a potential indicator of perceptually bound processing, which is more pronounced in younger adulthood and carries adverse implications not only for affective adjustment, but also for cognitive adaptation to novel environments. Finally, we characterise network communication profiles which link event segmentation processes across multiple timescales by providing perceptual and semantic scaffolding to context formation during perception, as well as during the subsequent episodic retrieval of these event representations.

## Materials & Correspondence

Correspondence and material requests should be addressed to R.P.

## Data statement

The raw data are available from https://camcan-archive.mrc-cbu.cam.ac.uk/dataaccess/ (Cam-Can) and https://db.humanconnectome.org/app/template/Login.vm;jsessionid=90091F006B15D5FA1D1D9C3ED2D465DD (HCP) upon completion of the relevant data use agreements.

## Code availability

We used already existing code, as specified in the main text.

## Conflict of interest

The authors declare no competing interests.

## Acknowledgments

K.S.G. and A.D.L. were funded by grants from the Medical Research Council (MR/N01233X/1) and a Wellcome Trust Strategic Award (104943/Z/14/Z). Data were provided by the Cambridge Centre for Ageing and Neuroscience (Principal Investigator: Lorraine K. Tyler; funders: the Biotechnology and Biological Sciences Research Council, the Medical Research Council Cognition & Brain Sciences Unit and the European Union Horizon 2020 LifeBrain project) and the Human Connectome Project, WU-Minn Consortium (Principal Investigators: David Van Essen and Kamil Ugurbil; 1U54MH091657; funders: the 16 NIH Institutes and Centers that support the NIH Blueprint for Neuroscience Research and the McDonnell Center for Systems Neuroscience at Washington University).

## Notes

### Competing Interest Statement

The authors have declared no competing interest.

